# Basal and stress-induced expression changes consistent with water loss reduction explain desiccation tolerance of natural *Drosophila melanogaster* populations

**DOI:** 10.1101/2022.03.21.485105

**Authors:** Vivien Horváth, Sara Guirao-Rico, Judit Salces-Ortiz, Gabriel E. Rech, Llewellyn Green, Eugenio Aprea, Mirco Rodeghiero, Gianfranco Anfora, Josefa González

**Author notes:** **Corresponding author:** Josefa González.

## Abstract

**Background:** Climate change is one of the main factors shaping the distribution and biodiversity of organisms, among others by greatly altering water availability, thus exposing species and ecosystems to harsh desiccation conditions. Insects are especially threatened by these challenging dry environments, because of their small size and thus large surface area to volume ratio. Integrating transcriptomics and physiology is key to advancing our knowledge on how species cope with desiccation stress, and these studies are still best accomplished in model organisms.

**Results:** Here, we characterized the natural variation of European *D. melanogaster* populations across climate zones and found that strains from arid regions were similar or more tolerant to desiccation compared with strains from temperate regions. Tolerant and sensitive strains differed not only in their transcriptomic response to stress but also in their basal expression levels. We further showed that gene expression changes in tolerant strains correlated with their physiological response to desiccation stress and with their cuticular hydrocarbon composition. Transposable elements, which are known to influence stress response across organisms, were not found to be enriched nearby differentially expressed genes. Finally, we identified several tRNA-derived small RNA fragments that differentially targeted genes in response to desiccation stress.

**Conclusions:** Our results showed that by integrating transcriptomics with physiological trait analysis we can pinpoint the genetic basis of the differences in tolerance to desiccation stress found in natural *D. melanogaster* populations. Moreover, we showed that, beyond starvation and aging, tRNA-derived small RNA fragments (tRFs) appear to be relevant post-transcriptional gene regulators in response to desiccation stress.

## BACKGROUND

Global climate changes, such as increased temperature and unpredictable changes in precipitation, pose a severe and widespread impact on organisms, from human health and food security, to species distribution and biodiversity (1–4). Of the natural disasters caused by climate change, droughts are among the costliest (5). These unpredictable patterns of precipitation are causing an increase in aridity and the expansion of drylands in many regions (6). Water related challenges are threatening many species, but insects are particularly vulnerable due to their small size and thus large surface area to volume ratio (7–9). In recent studies, a substantial (47-80%) decline in the abundance of some insect species was reported, partly due to climate change (10, 11). Because insects represent most of the animal diversity, and include many economically and ecologically extremely relevant species, such as bees, mosquitos and moths, understanding the adaptive responses of insects to climate change is crucial (12, 13).

Most of the insect-related climate change studies have so far focused on the effect of increased temperature (14–19), while response to desiccation conditions caused by changes in rainfall, humidity, and water availability have received less attention (12, 18, 20). *Drosophila* species are good models to study the physiological and genetic basis of adaptation to dry environments, as species of this genus have adapted to diverse climatic conditions during their recent evolutionary history, including arid regions (21–24). Indeed, geographical variation for desiccation tolerance among populations of several *Drosophila* species has been found (25–35). Among these species, *D. melanogaster* is an ideal model organism to further analyze desiccation stress response given its worldwide geographic distribution, the wealth of functional knowledge, and the genetic tools available (36).

There are several genome-wide studies investigating the underlying genetic architecture of desiccation tolerance in *D. melanogaster* (32, 37–40). However, knowledge on the genome-wide transcriptomic response is still limited, as most studies focused on the analysis of a few candidate genes in laboratory selected lines (41–46). The few transcriptomic studies available suggest that stress sensing, stress response, immunity, signaling and gene expression pathways are relevant for desiccation tolerance (38, 39, 44).

While the role of single nucleotide polymorphisms (SNPs) and chromosomal inversions in gene expression changes in response to desiccation have been investigated (38, 39), the potential role of transposable elements (TEs) as gene regulators in this stress response has not yet been studied. TEs are very powerful mutagens that can affect gene expression through a variety of molecular mechanisms (47, 48). Indeed, the adaptive role of TE insertions in response to stress conditions has been reported across organisms (49–51). Similarly, despite the growing evidence that points to tRNA-derived small RNA fragments (tRFs) as relevant post transcriptional gene regulators in stress response, their role in desiccation stress has not yet been studied either (52–55).

Finally, integrating physiology into the analysis of the genomic basis of desiccation stress response should help us better understand the underpinnings of this ecologically relevant trait. While three main physiological mechanisms have been related to desiccation tolerance in insects—water loss reduction, increased bulk water content, and water loss tolerance—the latter appears to be a less common mechanism in Drosophila (18, 56–59). Reduced water loss rate appears to be the most common mechanism to survive desiccation (22, 57, 59–62). Water loss happens mostly by two routes; the first occurs through the spiracles during the open phase in respiration (59, 63). The second is related to the cuticular hydrocarbons hindering transpiration (CHCs), which are the most prominent fatty acid-derived lipids on the insect body surface (58, 64). The variation in water loss through the cuticle has been related to the amount, chain length, and saturation of CHCs, with the most notable being a negative correlation between the length of the hydrocarbon chain and rates of water loss (8, 58, 60, 65). Finally, the role of increased bulk water content in desiccation tolerance is still not clear. While flies more tolerant to desiccation stress were found to have higher bulk water content (8, 35, 62, 66, 67), in other studies either no significant differences were described (68) or higher water content was associated with lower desiccation tolerance (69). To date, the physiological response to desiccation has been mainly studied in xeric Drosophila species or in *D. melanogaster* strains selected for desiccation tolerance. As such, a comprehensive picture in natural *D. melanogaster* populations is still not available (32–35, 70).

In this work, we assessed the natural variation for desiccation tolerance in European *D. melanogaster* natural strains collected in five climate zones (Figure 1A). We identified the transcriptomic responses and the underlying physiological traits that differentiate desiccation tolerant and sensitive strains. Finally, we also analysed the potential role of TEs and tRFs in this stress response.

**Figure 1.**
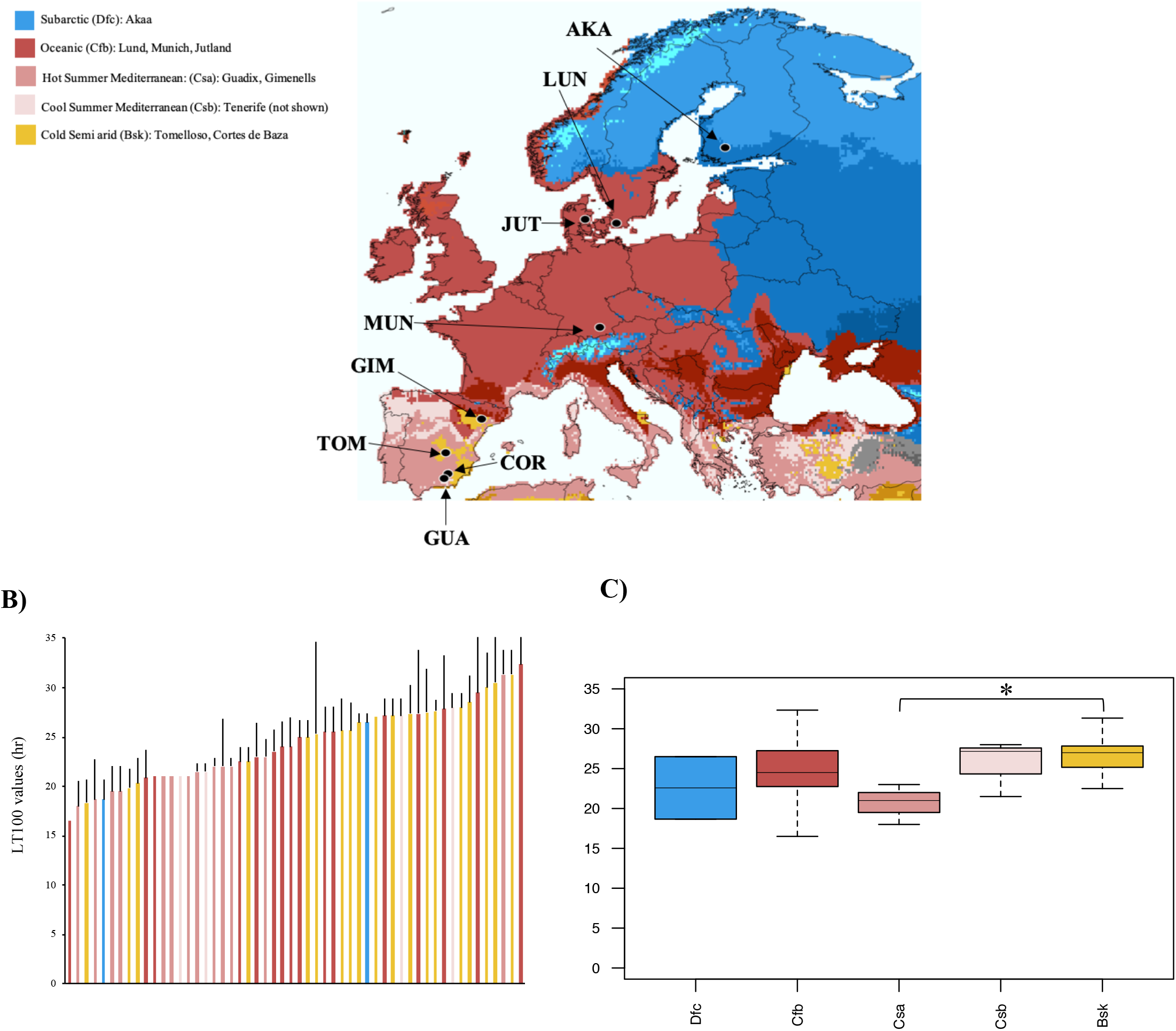
Natural variation in desiccation survival across European natural *D. melanogaster* populations. **A)** Geographical origin of the nine populations analyzed in this study. The location of the populations is indicated with arrows in a map of Europe coloured based on the Köppen-Geiger climate zones, except for Tenerife, which is not shown in the map. **B)** Desiccation survival of European natural populations. LT_100_ values for the 59 inbred strains that showed less than 10% control mortality are shown (name of strains can be found in Table S2A). The Y-axis represents the average hour when the flies in all the replicates were dead and the X-axis represents the individual strains coloured by the climate zone in which they were collected. Data are presented as mean values ± SD. **C)** Boxplot of the distribution of the LT_100_ values of the strains, grouped by climate zones. The boxplot shows median (the horizontal line in the box), 1st and 3rd quartiles (lower and upper bounds of box, respectively), minimum and maximum (lower and upper whiskers, respectively).

## RESULTS

### Altitude and evaporation correlate with desiccation tolerance in European natural *D. melanogaster* strains

To measure the variability in desiccation tolerance in *D. melanogaster* natural populations, we exposed 74 inbred strains from nine European locations to low humidity conditions (< 20% humidity) (Figure 1A and Table S1). These nine populations belong to five different climate zones including continental (Subarctic), temperate (Oceanic, Cool Summer Mediterranean, and Hot Summer Mediterranean climates) and dry (Cold Semi-arid) climates. LT_50_ values, representing the time when half the flies were dead, ranged from 12.5 to 29.7 hours, which is wider than those found in North American strains (17.9 to 20 hours (32); Figure S1A and Table S2A). LT_100_ values, representing the time when all the flies were dead, varied between 16.5 and 32.3 hours (Figure 1B and Table S2A). Similar differences in desiccation tolerance were found in populations from India collected from different latitudes, where the more tolerant strains survived about twice as long as the sensitive ones (LT_100_= 28.6 *vs.* 13.6) (34), while in Australian populations the LT_100_ variation was smaller (14.2 to 17.5 hours) (41). Thus, European populations showed similar or wider ranges of variation in survival times to desiccation stress compared with strains from other continents.

Flies from temperate climates have been shown to be more tolerant to desiccation stress than flies from tropical climates (32, 41, 57, 71). We thus tested whether there were significant differences in survival among flies from the five climates in our dataset (Figure 1). While no differences were found in the average LT_50_ values, we found significant differences in the average LT_100_ values (ANOVA, p-value = 0.467 and 0.023 for LT_50_ and LT_100_, respectively; Figure 1C and Table S2A). Pairwise comparisons showed significant differences between LT_100_ values of strains from the cold semi-arid (*Bsk*) and hot summer Mediterranean (*Csa*) climate zones: the cold semi-arid strains were more tolerant ((Tukey comparison, p-adj = 0.01; Figure 1C and Table S2A).

Finally, desiccation tolerance has been correlated with altitude, latitude, and with environmental variables such as annual precipitation and minimum temperature (20, 32, 34, 72). We did not find a significant correlation between latitude, longitude or altitude and the desiccation tolerance of the strains (LT_100_; linear model, p-value = 0.648 for altitude, p-value = 0.853 for latitude, and p-value = 0.686 for longitude; Table S2A). On the other hand, we found that the LT_100_ values significantly correlated with the interaction of altitude and evaporation (p-value = 0.0005, adjusted R-squared: 0.135; Table S2A; see Methods).

### Desiccation tolerant and sensitive strains differed in the number, the direction of expression change, and the function of genes that respond to stress

To investigate the transcriptional response to desiccation stress, we generated whole female RNA-seq data for three tolerant and three sensitive strains chosen from the extremes of the LT_50_ distribution (Figure S1B, Table S2B; see Methods). We used *Transcriptogramer,* which takes into account protein-protein interactions to perform differential expression of functionally associated genes (clusters) (73), to investigate the overall response to desiccation stress, by analyzing the six strains together (“All DEGs”). We also investigated whether tolerant (“Tolerant DEGs”) and sensitive (“Sensitive DEGs”) strains differed in their transcriptomic response to desiccation stress.

When analyzing the six strains together, we identified five clusters of DEGs with 92% (1292 out of 1405) of the genes down-regulated (Figure 2A and Table S3A). Genes in the two clusters with the biggest number of down-regulated genes were enriched for metabolic processes (RNA, nitrogen compounds, macromolecules) and gene expression functions (Figure 2A and Table S3A). In the other two down-regulated clusters, genes were enriched for chitin metabolic process, fatty acid biosynthetic process, and cuticle development (Figure 2A and Table S3A). Note that genes related to gene expression, RNA metabolism, and chitin metabolism have been previously associated with desiccation stress response (28, 38, 39). On the other hand, up-regulated genes were enriched for cell communication, signaling, and response to stimulus functions (Figure 2A and Table S3A). Response to stimulus and environmental sensing have also been previously associated to desiccation stress response in *D. melanogaster* and *D. mojavensis* (28, 39).

**Figure 2.**
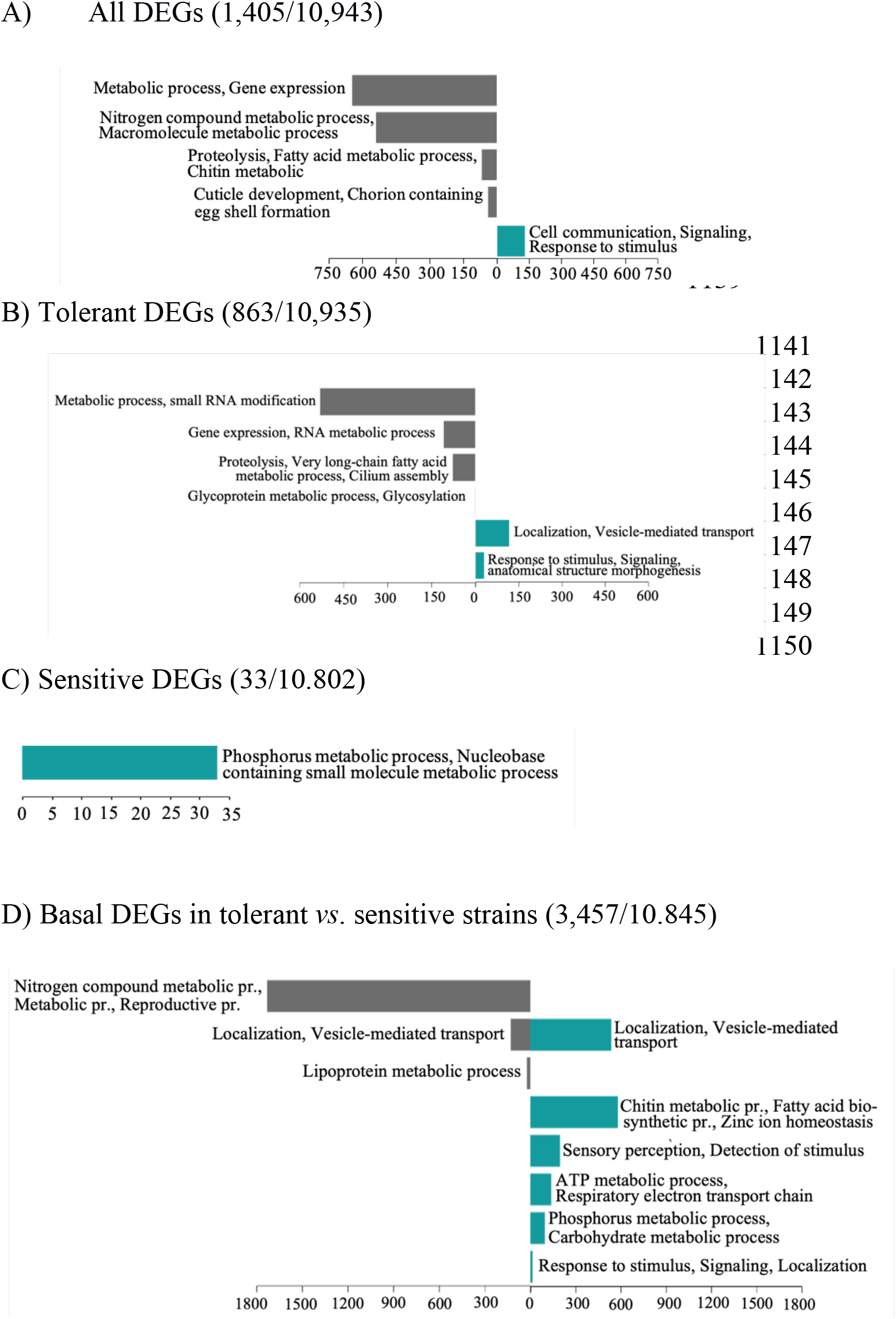
Biological process GO enrichment of the differentially expressed gene clusters identified by *Transcriptogramer*. A) GO enrichment of the DEGs when all six strains are analysed together. B) GO enrichment of the DEGs in tolerant strains. C) GO enrichment of the DEGs in sensitive strains. D) GO enrichment of the DEGs in basal conditions when comparing tolerant vs sensitive strains. In all the plots, the X-axis represents the number of genes contained in the clusters. Grey indicates enrichment for down-regulated genes, while blue indicated enrichment for the up-regulated genes. In parenthesis the number of DEGs/number of genes analysed are given.

The same pattern was found in the “Tolerant DEGs” with 83% of the genes down-regulated (716 out of 863 Figure 2B and Table S3B). This result is not unexpected, as 86% of the “Tolerant DEGs” overlap with the “All DEGs”. The four down-regulated and two up-regulated clusters were enriched for similar GO terms as the ones found in the “All DEGs” analysis, with the up-regulated clusters also enriched for localization and transport functions (Figure 2B and Table S3B).

On the other hand, a very small number of genes (33 genes) were differentially expressed in sensitive strains, and they were all up-regulated and enriched for nucleotide metabolic and catabolic processes (Figure 2C and Table S3C). This low number of genes suggested that sensitive strains have a limited coordinated desiccation stress response.

We found a considerable overlap among our DEGs, considering “All DEGs”, “Tolerant DEGs” and “Sensitive DEGs”, and genes previously related to desiccation stress response (379 out of 1,524 DEGs (25%); Table S4A), including four DEGs that affect the cuticular composition of *D. melanogaster* (74), and 10 DEGs that overlap with the core set of candidate genes identified in the cross-study comparison carried out by Telonis-Scott *et al.* (2016) (Table S4B) (39). While we found overlap with previously described candidates, our transcriptomic analysis also identified new candidate genes (see below).

Overall, tolerant and sensitive strains differed in their transcriptomic response to stress not only in the number of desiccation-responsive genes but also in the direction of the change of expression and in the gene functions (Figure 2A-C).

### Tolerant strains have a higher level of basal expression of desiccation-responsive genes

Basal transcriptional states have been associated with differential response to cold stress and bacterial infection (62, 75). We thus compared the gene expression in tolerant *versus* sensitive strains in basal conditions. We found eight gene clusters containing 3,457 DEGs, which included 851 genes that have been previously related to desiccation stress; eight of them affecting the cuticular composition of *D. melanogaster* (Table S3D-S4A). Moreover, 15 of the DEGs in basal conditions overlap with the core set of candidate genes identified in the cross-study comparison of Telonis-Scott *et al*. (2016) (Table S4B) (39).

Down-regulated gene clusters were mostly enriched in metabolic processes, while up-regulated clusters contained genes related to response to stimulus, ion transport, sensory perception, cell communication, metabolic processes such as chitin metabolic processes, fatty acid elongation and very long chain fatty acid biosynthetic process (Figure 2D and Table S3D).

Overall, our results showed that tolerant and sensitive strains differed in their basal gene expression levels, with tolerant strains showing higher levels of basal expression of genes previously associated with desiccation tolerance such as response to stimulus, chitin metabolic process, and fatty acid elongation (Figure 2D and Table S3D).

### Desiccation-responsive genes are enriched for highly expressed genes in the ovary

We tested whether DEGs in response stress, and DEGs when comparing tolerant and sensitive strains in basal conditions were enriched for highly expressed genes in the ovary, as has been previously described (37). Using the *Drosophila Gene Expression Tool* (DGET) (76), we found that stress-response DEGs, including hub DEGs (see Methods), were enriched for highly expressed genes in the ovary (Hypergeometric test, p-value < 0.0001, for all comparisons; Table S5A-C and Table S6). On the other hand, basal DEGs were enriched for highly expressed genes in head, digestive system, carcass and ovary (Hypergeometric test, p-value < 0.0001, for all comparisons). However, basal hub DEGs were enriched for highly expressed genes in the ovary and digestive system (Hypergeometric test, p-value < 0.0001, for all comparisons, Table S5D-S6).

### *nclb, Nsun2*, and *DBp73D* genes affect desiccation tolerance in *D. melanogaster*

Besides detecting genes previously known to play a role in desiccation tolerance, our transcriptomic analysis also identified new candidate genes (Table S3). We chose three hub genes among the ones with the highest Maximal Clique Centrality (MCC) values, *nclb, Nsun2, Dbp73D,* to perform functional validation experiments (Table S6; see Methods; 77). These genes were (i) mostly expressed in the ovary and digestive system; (ii) related to gene expression and RNA methylation (Table 1); and (iii) were down-regulated in the “All DEGs” and “Tolerant DEGs” groups (Table S3). In the case of *nclb*, the insertional mutant and the two reciprocal crosses of the RNAi transgenic line analyzed showed a lower level of expression and higher survival in desiccation stress conditions when compared with background strains (Log-rank test, p-value < 0.0001 in all three comparisons; Figure 3, Table 1, and Table S7). These results are consistent with the observed down-regulation of the *nclb* gene in tolerant strains in desiccation stress conditions (Table S3).

**Figure 3.**
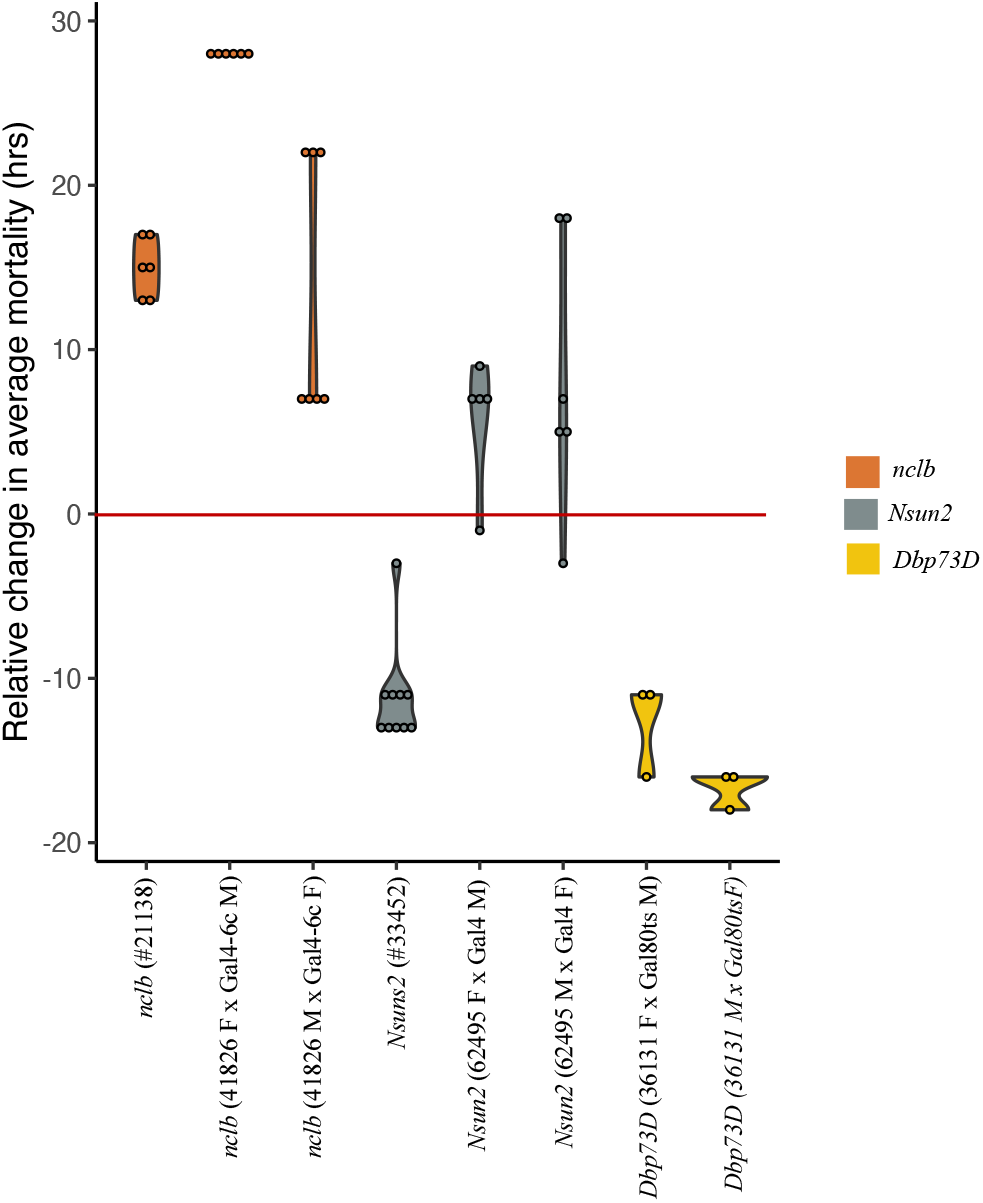
Functional validation of *nclb, Nsun2* and *Dbp73D* genes. Relative change in average mortality at the end of the desiccation assay comparing gene disruption and knock-down strains for the three candidate desiccation-responsive genes with control strains.

**Table 1.** Functional validation of three candidate desiccation-responsive genes. Expression level and survival of gene disruption and RNAi reciprocal crosses relative to the control strains (F= female and M= male).

| Gene name/<br>Flybase ID | Gene function | Gene disruption/RNAi (stock number) | qRT-PCR results (p-value) | Survival of the gene disruption /RNAi strain | Log-rank test p-value |
| --- | --- | --- | --- | --- | --- |
| *nclb*/<br>FBgn0263510 | Regulation of gene expression | *P-element* insertion in first intron (#21138) | Lower expression (3.80E-03) | Higher survival | <0.0001 |
|  |  | RNAi (#41826) F x Gal4-6c M | Lower expression (0.03) | Higher survival | <0.0001 |
|  |  | RNAi (#41826) M x Gal4-6c F | Marginally lower expression (0.067) | Higher survival | <0.0001 |
| *Nsun2*/<br>FBgn0026079 | RNA methylation, tRNA methylation | *P-element* insertion in first intron (#33452) | Lower expression (0.024) | Lower survival | <0.0001 |
|  |  | RNAi (#62495) F x Gal4 M | Lower expression (5.07E-04) | Higher survival | <0.0001 |
|  |  | RNAi (#62495) M x Gal4 F | Lower expression (0.03) | Higher survival | <0.0001 |
| *Dbp73D*/<br>FBgn0004556 | RNA binding | RNAi (#36131) F x Gal80ts M | Lower expression (0.007) | Lower survival | <0.0001 |
|  |  | RNAi (#36131) M x Gal80ts F | Lower expression (0.01) | Lower survival | <0.0001 |

For *Nsun2*, we found that while the expression of the gene in the insertional mutant and the two reciprocal crosses of the RNAi transgenic line was lower, the survival to desiccation stress was lower for the insertional mutant but higher for the RNAi reciprocal crosses (Log-rank test, p-value: <0.0001 for the three comparisons; Figure 3, Table 1 and Table S7). These results are consistent with a role of *Nsun2* in desiccation tolerance and suggest that the effect of this gene is background dependent. Finally, for *Dbp73D*, the two reciprocal crosses performed with the RNAi line showed lower gene expression and lower survival under desiccation stress conditions (Log-rank test, p-value: <0.0001; Figure 3, Table 1, and Table S7). This result suggests that the effect of *Dbp73D* on desiccation tolerance is also background dependent as we found this gene to be down-regulated in tolerant strains. Overall, we found that all three genes affect the desiccation survival of the flies, however in some cases this effect depends on the genetic background.

### Differentially expressed genes in response to desiccation are not enriched for TE insertions

TEs have previously been shown to affect gene expression in response to stress in *D. melanogaster* (*e.g*. 78, 79). As a first step towards the analysis of the role of TE insertions in desiccation stress response, we tested whether DEGs in response to desiccation stress and DEGs when comparing tolerant *vs*. sensitive strains basal expression were enriched for TE insertions. We used the *de novo* TE annotations for the six genomes analyzed to identify TEs located either inside genes or in the 1kb upstream and downstream gene regions (80). We found that while 3% to 9.8% of the DEGs in response to desiccation were located nearby a TE insertion, this percentage was not significantly higher than expected when compared with the genome-wide distribution of genes nearby TEs (Table 2 and Table S8). Similar results were obtained for basal DEGs, where 12% of them were located nearby TEs, but this percentage was indeed smaller than expected (Table 2 and Table S8). Although DEGs were not significantly enriched for nearby TE insertions, we cannot discard that some of the identified TEs located nearby DEGs could be responsible for the differences in expression detected. Additional functional validation experiments would be needed to test this hypothesis.

**Table 2.**
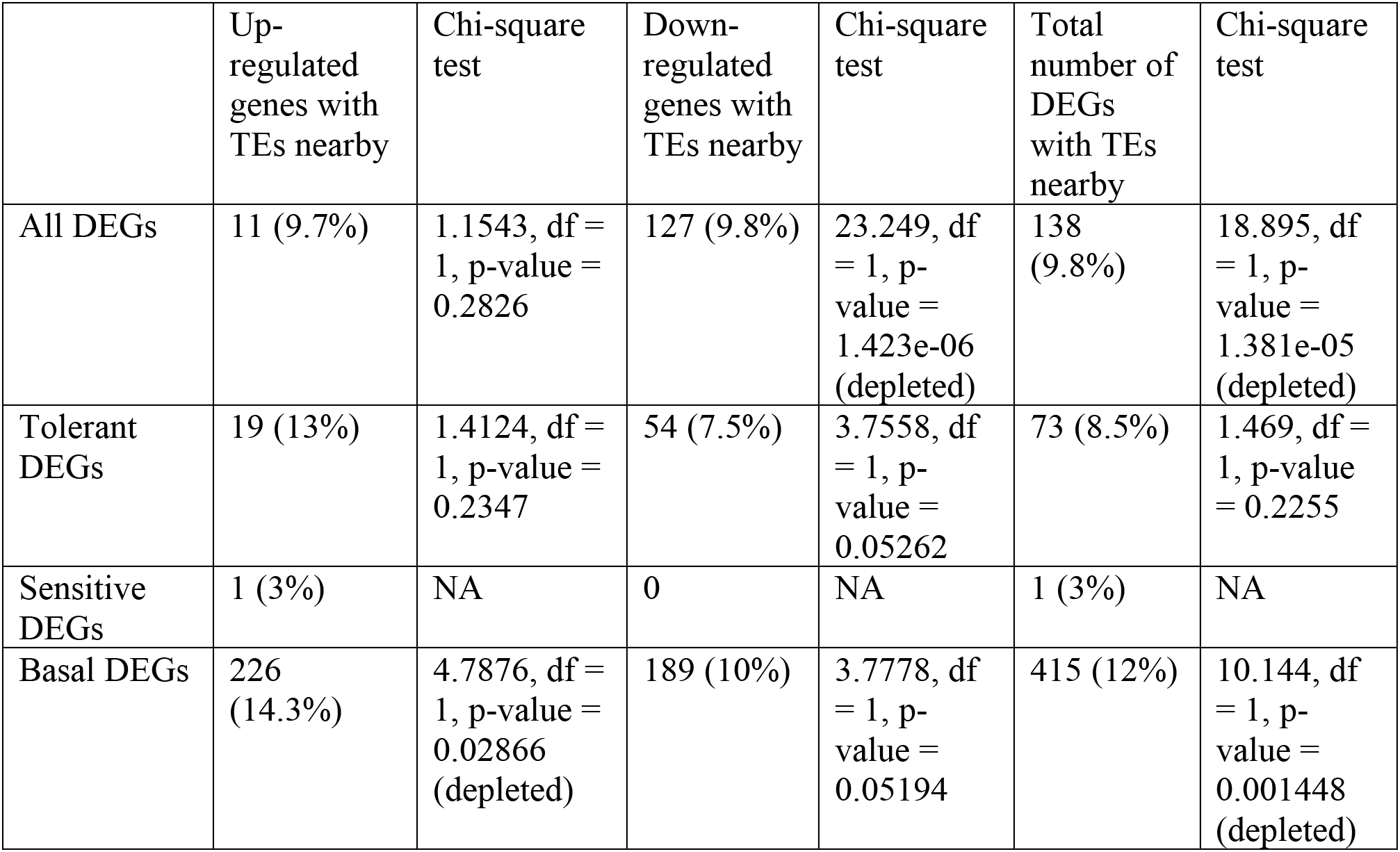
Number of differentially expressed genes located nearby transposable element (TE) insertions. d.f.: degrees of freedom

### tRNA derived fragments differentially target genes in response to desiccation stress

While tRNA-derived small RNAs (tRFs) are known to inhibit the translation of protein coding genes during starvation stress and aging, due to complementary sequence matching, their potential role in desiccation stress response has not yet been assessed (52, 53). To test whether tRFs could play a role in desiccation stress response, we sequenced whole female small RNAs under control and desiccation stress conditions in the tolerant and in the sensitive strains. We identified the genes targeted by tRFs in tolerant and sensitive strains and focused on those genes that were differentially targeted in response to desiccation stress *i.e.* genes that gain or lose targeting (Table S9; see Methods). For tolerant strains, 106 genes were targeted by tRFs in response to stress while 332 genes lost targeting in response to stress (Table S9A). While the same number of genes were targeted by tRFs in response to stress in sensitive strains, the number of genes that lost targeting was smaller (53 genes; Table S9A). The majority of genes that lost targeting in response to stress overlap between sensitive and tolerant strains (87%: 46 out of 53). This overlap is much smaller for the genes that are targeted in response to stress suggesting that tRFs target different genes in tolerant and sensitive strains (37%: 39 out of 106).

We tested whether DEGs in tolerant and sensitive strains overlapped with genes that were post-transcriptionally targeted by tRFs (Table S9B). The overlap ranged from 1% to 11% suggesting that different sets of genes are controlled at the transcriptional and post-transcriptional levels. The overlap at the biological process level was also small, suggesting that transcriptional and post-transcriptional control of desiccation responsive genes is different both at the gene and at the biological process level (Figure 2, Table 3, and Table S9C). We finally tested whether the genes that were differentially targeted by tRFs in response to desiccation stress in tolerant and sensitive strains have been previously identified as desiccation-responsive genes (Table S9D). Up to 5% (40/423) of genes targeted by tRFs have previously been identified as desiccation-responsive genes (Table S9D).

**Table 3.** GO enrichment analysis of the genes that gain or lose targeting by tRFs in response to desiccation stress in tolerant and in sensitive strains.

|  | **Targeting appears in response to desiccation stress** | **Enrichment Score** | **Targeting disappears in response to desiccation stress** | **Enrichment Score** |
| --- | --- | --- | --- | --- |
| **Tolerant strains** | Response to stimulus | 1.7 | Nervous system development | 3.9 |
|  |  |  | Regulation of developmental process, neurogenesis | 2.8 |
|  |  |  | Regulation of metabolic process | 2.4 |
|  |  |  | Response to stimulus | 2 |
|  |  |  | Regulation of signaling | 1.9 |
| **Sensitive strains** | Nervous system development | 1.9 | NS | NA |
|  | Signaling, response to stimulus | 1.4 |  |  |
|  | Cell adhesion | 1.4 |  |  |
|  | Response to growth factor | 1.3 |  |  |

Overall, these results suggest that tRFs do play a role in desiccation stress response, with the majority of tRFs targeting different genes in tolerant and sensitive strains, and allow the identification of new candidate desiccation-responsive genes.

### Differences in the cuticular hydrocarbons and in respiration rate are associated with reduced water loss in desiccation tolerant strains

Besides investigating the transcriptional response to stress and the potential role of TEs and tRFs in this response, we also characterized the physiological response to desiccation in tolerant and sensitive strains. We measured three physiological traits that have been associated with the level of desiccation tolerance in flies—water content, rate of water loss and respiration rate—and we characterized the cuticular hydrocarbon composition of tolerant and sensitive strains (34, 35, 60, 69).

#### Tolerant strains have lower bulk water content

We first checked the bulk water content in the 10 most tolerant and 10 most sensitive strains of the LT_50_ distribution (Figure S1A; see Methods). We found that the tolerant strains had significantly lower initial water content compared to the sensitive strains (45% *vs*. 50% on average, Wilcoxon test, p-value: < 0.0001, Figure 4A and Table S10A), which is consistent with previous studies (69).

**Figure 4.**
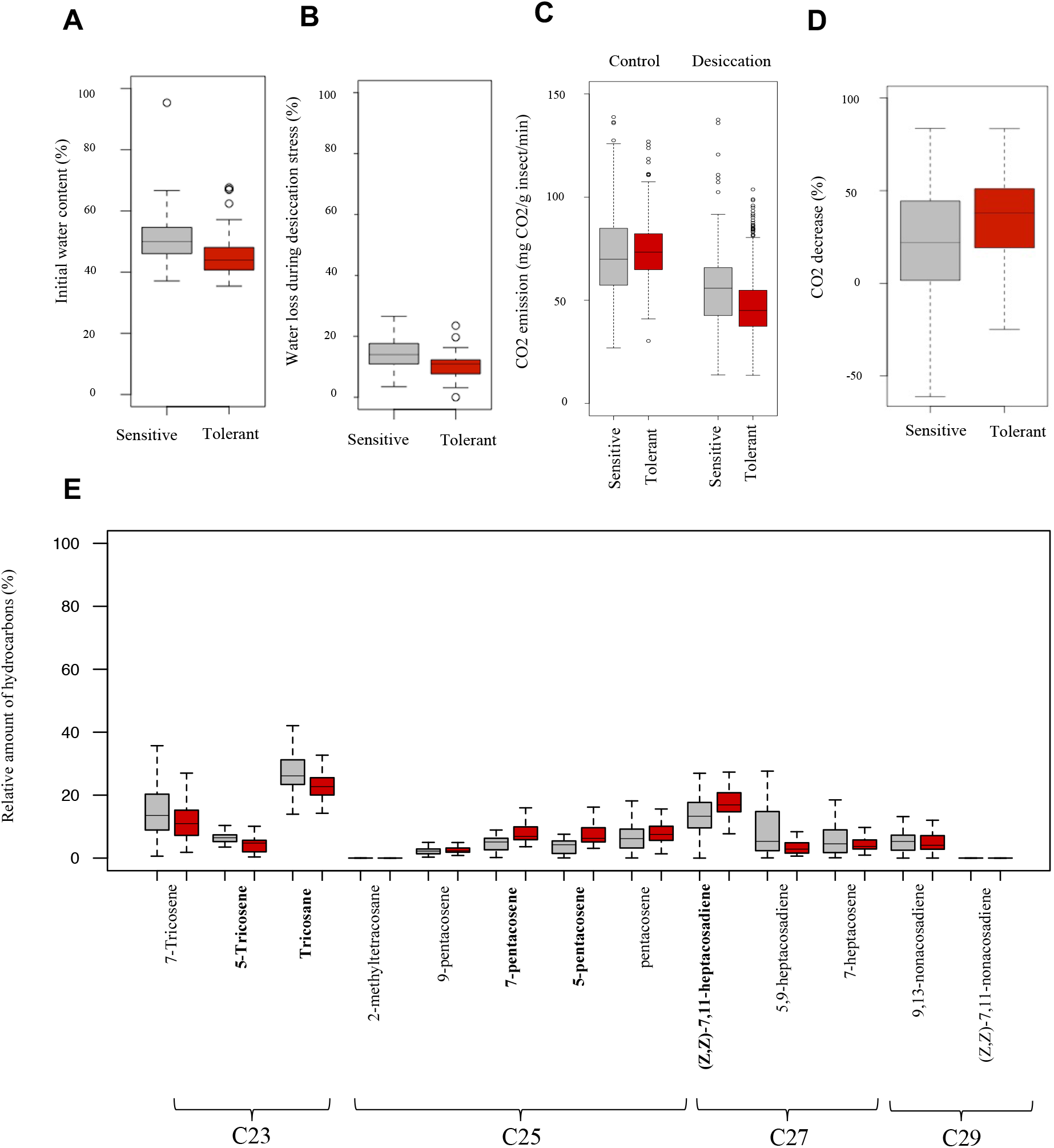
Desiccation-related physiological traits and cuticular hydrocarbon variation in natural European populations. Results of **A)** initial water content in sensitive and tolerant strains. **B)** percentage of water loss during desiccation stress in sensitive and tolerant strains. **C)** Respiration rate under control and desiccation stress conditions in sensitive and tolerant strains. **D)** Percentage of CO_2_ decrease (respiration) in response to desiccation stress in sensitive and tolerant strains. **E)** Relative amount of hydrocarbons in sensitive (grey) and tolerant (red) strains. Hydrocarbons that showed significant differences between sensitive and tolerant strains are depicted in bold. All boxplots show the median (the horizontal line in the box), 1st and 3rd quartiles (lower and upper bounds of box, respectively), minimum and maximum (lower and upper whiskers, respectively).

#### Tolerant strains lose less water during desiccation stress compared with sensitive strains

Next, we quantified the amount of water loss by measuring the fly’s weight before and after three hours of desiccation stress. We found that sensitive strains lose on average 15% of their water content while the tolerant strains lose 10% on average (Wilcoxon, p-value < 0.0001; Figure 4B and Table S10B). These results are consistent with previous studies performed both in populations selected for desiccation stress tolerance and natural populations (34, 62).

#### Tolerant strains decrease their respiration rate more in desiccation stress conditions

We measured the respiration rate of tolerant and sensitive strains in control and desiccation stress conditions. We compared GLMM with and without interaction between the experimental condition (control and desiccation stress) and the phenotype of the strains (tolerant and sensitive), and we found that the model including the interaction fitted the data better (LRT, p-value < 0.0001) (Table S11). Tolerant strains have a higher respiration rate in control conditions compared with sensitive strains (Figure 4C). However, after desiccation stress, the sensitive strains lower their respiration rate by 23% on average while the tolerant strains do so by 35% (Figure 4D).

#### Tolerant strains have higher relative amount of desaturated hydrocarbons

The level of cuticular transpiration, which is another influential factor in water loss, depends on the composition of the cuticle. Thus, we next analyzed the cuticular hydrocarbon (CHC) composition in tolerant and sensitive strains in control conditions (see Methods). Overall, we identified 13 main hydrocarbons with chain lengths varying between 23 and 29 carbons, including three saturated (n-alkanes) and ten desaturated compounds (alkene, alkadiene) (Figure 4E, Table 4, Table S12A). We performed principal component analysis (PCA) to explore the variability of the strains in terms of CHC composition. We found that although tolerant and sensitive strains differed in the relative amounts of individual hydrocarbons (see below), neither PC1 nor PC2 separated the 10 tolerant from the 10 sensitive strains when considering the total CHC composition (Figure S2A). However, PC1 did separate the three tolerant from the three sensitive strains for which we characterized the transcriptomic response to desiccation stress and explained 63.72% of the variation (Figure S2B). Note that these six strains are on the extremes of the phenotypic distribution for water loss measurements (Table S10B).

**Table 4.** Cuticular hydrocarbons (CHCs) identified in the 10 most tolerant and 10 most sensitive strains. CHCs with a statistically significant different amount between the tolerant and sensitive strains are marked in bold. Note that, linear retention index denotes the behavior of compounds on a gas chromatograph according to a uniform scale determined by the internal standard (127).

| Linear retention index | Component | Formula (acronym) | Hydrocarbon type | Saturated/unsaturated |
| --- | --- | --- | --- | --- |
| 2279 | 7-Tricosene | C23H46 (7-C23:1) | Alkene | Desaturated |
| 2290 | **5-Tricosene** | C23H46 (5-C23:1) | Alkene | Desaturated |
| 2298 | **Tricosane** | C23H48 (n-C23) | Alkane | Saturated |
| 2464 | 2-Methyltetracosane | C25H52 (2-Me-C25) | Alkane | Saturated |
| 2473 | 9-pentacosene | C25H50 (9-C25:1) | Alkene | Desaturated |
| 2480 | **7-pentacosene** | C25H50 (7-C25:1) | Alkene | Desaturated |
| 2480 | **5-pentacosene** | C25H50 (5-C25:1) | Alkene | Desaturated |
| 2498 | pentacosane | C25H52 (n-C25) | Alkane | Saturated |
| 2657 | **(Z,Z)-7,11 heptacosadiene** | C27H52 (7,11-C27:2) | Alkadiene | Desaturated |
| 2670 | 5,9-heptacosadiene | C27H52 (5,9-C27:2) | Alkadiene | Desaturated |
| 2693 | 7-heptacosene | C27H54 (7-C27:1) | Alkene | Desaturated |
| 2841 | 9,13-nonacosadiene | C29H56 (9,13-C29:2) | Alkadiene | Desaturated |
| 2872 | **(Z,Z)-7,11-nonacosadiene** | C29H56 (7,11-C29:2) | Alkadiene | Desaturated |

Tolerant strains had a higher relative amount of desaturated hydrocarbons (Wilcoxon test, p-value = 0.004), and a higher desaturated:saturated balance ratio compared with sensitive strains, as previously reported (Wilcoxon test, p-value = 0.004191; Table S12A; (69). However, the percentage of 7,11:Cn alkadienes, which was previously reported to be negatively correlated with desiccation tolerance, was found to be slightly positively correlated in our strains (Spearman’s correlation = 0.203, p-value = 0.02; (70). The relative percentage of longer chain hydrocarbons (≥ 27C) was not correlated with desiccation tolerance (Spearman’s correlation = −0.19, p-value = 0.828; Table S12A; (70). However, in Foley & Telonis-Scott (2011) hydrocarbons with ≥ 27 carbons represented approximately half of the total CHC composition; while in our European strains, compounds with ≥25 carbons or more are the most abundant. This is likely explained by the different latitudes in which populations from these two studies were collected (33). If we focus on ≥25C compounds, we did find a higher percentage in the tolerant compared with sensitive strains (Wilcoxon test, p-value < 0.0001). Moreover, the percentage of ≥25C compounds positively correlated with the LT_100_ (Spearman’s correlation = 0.178, p-value = 0.041) (Table S12A). In line with these results, we also found that tolerant strains had a higher relative percentage of 5-pentacosene, 7-pentacosene and 7,11-heptacosadiene, which are all compounds with ≥25C (Wilcoxon, p-value < 0.0001 in all cases; Fig 4D); while sensitive strains had a higher relative percentage of 5-tricosene and tricosene (Wilcoxon, p-value:<0.0001 in all cases), which are compounds with 23C (Figure 4E).

Overall, tolerant and sensitive strains at the extremes of the phenotypic distribution differed in the global CHC composition, and all the strains tested differed in the relative percentage of individual CHCs. While we did not find some of the previously described correlations between particular CHC and desiccation tolerance, this is most likely explained by the different CHC composition of the European strains in which CHC with 25 or more carbons, instead of CHC with 27C or more carbons, represent approximately half of the total CHCs.

## DISCUSSION

In this study, we characterized the variation in desiccation tolerance of natural European *D. melanogaster* strains from temperate, continental, and arid regions (Figure 1). While in previous studies in Australia (41, 57), South America (71), and North America (32) temperate strains were found to be more tolerant compared to tropical ones, here we showed that in Europe, strains from arid regions are similar or more tolerant to desiccation compared to strains from temperate regions (Figure 1C). We also found that variation in desiccation resistance in European strains can be partly explained by altitude and evaporation. The importance of altitude was previously shown in Indian populations of *D. melanogaster*, where flies from highlands were more tolerant compared to flies from lowlands (34).

Besides characterizing the natural variation in desiccation stress tolerance, we sought to uncover the physiological traits that influence this variation and the coordinated response of genes which orchestrate it. In control conditions, the tolerant strains showed a higher level of respiration rate (Figure 4D and Table S11). Consistent with this result, genes related to respiration *i.e.* respiratory electron transport chain, and cellular respiration, were up-regulated in tolerant strains in control conditions (Figure 2D and Table S3D). Genes related to ion transport were also up-regulated in the tolerant strains compared to the sensitive ones in control conditions (Table S3D). Ion homeostasis genes have been suggested to be involved in water retention by the Malpighian tubules and other cells throughout the body (32, 38), and have been related to desiccation survival before by regulating water retention in the flies (81).

Although tolerant strains have a higher respiration rate in control conditions, following desiccation stress, they lower their respiration rate further than the sensitive strains (Figure 4D), and were consistently found to lose less water (Figure 4B). Reduced water loss after desiccation stress has previously been found in desiccation tolerant *D. melanogaster* strains and xeric *Drosophila* species (8, 22, 34, 35, 62, 68). Moreover, we found that genes related to metabolic processes were down-regulated after desiccation stress in the tolerant strains, thus probably causing a lowered metabolism (Figure 2B). Reduction in the metabolic rate is known to reduce the need to open the spiracles, which is consistent with tolerant flies losing less water (82–84).

We found that besides differences in respiration rate, differences in cuticular hydrocarbons (CHC) composition between tolerant and sensitive strains are also likely to contribute to reduced water loss in tolerant strains (Figure 4E). Moreover, genes related to very long chain fatty acid elongation (bigger than 20C; 85) and chitin metabolic process were up-regulated in the tolerant strains under basal conditions (Figure 2D). However, they were down-regulated after desiccation stress (Figure 2B). This result suggests that the tolerant strains do not improve the water retaining properties of the cuticle during desiccation stress by increasing the production of cuticular proteins as has been suggested in *D. mojavensis,* where genes involved in chitin metabolism and cuticle constituents were found to be up-regulated after desiccation stress (28). Thus, it seems that the tolerant strains have a different, more favourable CHC composition under basal conditions compared with sensitive strains which might be related with their capacity to better survive low humidity conditions.

Tolerant strains also showed an increased expression of genes related to response to stimulus, signaling, localization and transport after desiccation stress (Figure 2B). Furthermore, tolerant strains also up-regulated genes related to sensory perception and detection of chemical stimulus under basal conditions compared to sensitive strains (Figure 2D). These results suggested that the response to desiccation has an important environmental sensing component, as has been previously suggested in *D. melanogaster* and in *D. mojavensis* (28, 38, 39). Indeed *Pkd2* (*Polycystic kidney disease 2*), *pyx* (*pyrexia*), and *pain* (*painless*) genes involved in hygroreception, a sense that allows the flies to detect changing levels of moisture in the air, were found to be differentially expressed in tolerant strains under basal conditions (Table S3D; 38, 39). Moreover, 27% (14 out of 52) of odorant binding proteins (*Obp*) that are also associated with hygroreception (46), were up-regulated in tolerant strains in basal conditions, which also underpins the importance of environmental sensing in control conditions (Table S3D).

The transcriptomics characterization of desiccation tolerant and sensitive strains also allowed us to pinpoint new desiccation-responsive gene candidates. Two of the three desiccation-responsive genes validated in this work, *Dbp73D* and *Nsun2*, have already been shown to be important in stress responses in *Arabidopsis thaliana* and mice (86, 87). In *Arabidopsis thaliana, Dbp73D* mutants show increased tolerance to salt, osmotic stress and heat stress (87). In mice, the *Nsun2* gene has been shown to be repressed amid oxidative stress, and the loss of *Nsun2* gene altered the tRF profiles in response to oxidative stress (86). In Drosophila, *Nsun2* mutants have been shown to cause loss of methylation at tRNA sites (88). These results suggest that *Dbp73D* and *Nsun2* appear to be involved in general stress response across organisms and that tRFs are part of the stress response. In this work, we also investigated the potential role of tRFs in desiccation stress response and found tRFs that differentially target genes in tolerant and sensitive strains in response to desiccation stress (Table 3). Previous works in *D. melanogaster* pinpoint the role of tRFs in cellular starvation response through the regulation of translation of both specific and general mRNAs (53). We found that the overlap between DEGs and genes targeted by tRFs was small suggesting that different genes are controlled at the transcriptional and post-transcriptional levels. However, further functional validation analysis is needed to confirm the role in desiccation stress response of the genes targeted by tRFs.

## CONCLUSIONS

Overall, we found that *D. melanogaster* natural populations from arid regions are similar or more tolerant to desiccation stress compared with populations from temperate regions, and that this tolerance correlates with altitude and evaporation. Combining gene expression profiling with physiological trait analysis allowed us to pinpoint the genetic basis of the physiological response to desiccation stress. We found that differences in both basal gene expression and stress response are consistent with differences in the cuticular hydrocarbon composition and in the respiration rate, which altogether contribute to explain the water loss reduction of tolerant strains. Transcriptomic analyses also allowed us to identify new candidate desiccation-responsive genes, three of which were functionally validated. Finally, we identified genes that are differentially targeted by tRFs in response to stress, suggesting that tRFs previously involved in starvation resistance and aging could also play a role in desiccation stress response in *D. melanogaster*. The results obtained in this work can also have considerable practical implications, for example by helping to understand the potential for spread, adaptation and damage of economically important species in the light of ongoing climate change. An example might be the case of an invasive alien species belonging to the same genus, *Drosophila suzukii,* one of the major agricultural pests worldwide (89, 90).

## METHODS

### Fly husbandry

Flies were collected in 2015 from nine European locations by members of the *DrosEU* consortium (Figure 1 and Table S1). These nine locations belong to five different climate zones according to the Köppen-Geiger climate classification: Subarctic (*Dfc*), Oceanic (*Cfb*), Cool Summer Mediterranean (*Csb*), Cold Semi-arid (*Bsk*) and Hot Summer Mediterranean (*Csa*) (Table S1; 50, 91). In total, 74 inbred strains were generated from the aforementioned natural populations. Flies were inbred for 20 generations, except for two strains which were inbred for 21 generations and 11 strains that were too weak to continue with the inbreeding process and were thus stopped before reaching 20 generations (Table S1). Fly stocks were kept in vials at 25 °C and 65% relative humidity with 12h day and night cycles.

### Desiccation survival experiments

For each of the 74 strains analyzed, three replicates of 15 individuals each (except for 63 replicas with 10-18 individuals) were performed for desiccation stress conditions and 3 replicas (except for two strains where 1-2 replicas were performed) of 15 individuals each (except for 4 replicates that were performed with 9-10 flies) were performed for control conditions (Table S2A). In both conditions, 4-8 day-old females were used. Also in both conditions, the vials were closed with cotton and sealed with parafilm to stop the airflow. In treated conditions, flies were put in empty vials and three grams of silica gel (Merck) were placed between the cotton and the parafilm, so they were starved and desiccated. Control vials were prepared similarly, except that they contained 1mL of 1% agar on the bottom of the vial to prevent desiccation (agar provides hydration but is not a food source; (43). Temperature and humidity were continuously monitored using three *iButtons* (Mouser electronics) (Table S2). Fly survival was monitored every four hours until hour 12, and at shorter intervals afterwards (1 to 3.5 hours). Flies that died before the first survival check were considered to have been injured during the experiment setup and were not included in the analysis (Table S2). For 15 out of the 74 strains, >10% mortality was observed in control conditions and thus were not further analyzed (Table S2A). Lethal time 50 (LT_50_, the time when 50% of the flies are dead) was calculated using *Probit* analysis (92) for all the strains (59 strains). LT_100_ was calculated for all the strains for which mortality data were collected until the end of the experiment (54 strains).

### Correlation of LT_100_ with environmental and geographical variables

To test if the LT_100_ data followed a normal distribution, we used the Shapiro-Wilk normality test. As the data were normally distributed, we performed an ANOVA analysis to test if there were differences in the average of the LT_100_ values among different climate zones, followed by a Tukey Test to check which climates differ from each other. We then tested if the LT_100_ was correlated with geographical or environmental variables. The geographical variables used were longitude, latitude and altitude (Table S1). For environmental data, we used two different sources: (i) *WorldClim* (www.worldclim.org; (93) and (ii) Copernicus (*ERA5*) (Table S1) (94). Environmental variables related to temperature and precipitation have been shown to explain the greatest variability in desiccation resistance in Drosophila (20); so we used the 19 bioclimatic variables from WorldClim, which were derived from the monthly temperature and rainfall values between 1970-2000, and the yearly maximum and minimum temperature. We used the R package *raster* (v. 2.6-7) for downloading these data (95). We also used evaporation (the accumulated amount of water that has evaporated from the Earth’ surface, including a simplified representation of transpiration (from vegetation), into vapor in the air above) and solar radiation (clear-sky direct solar radiation at surface) data from the year previous to the collection date obtained from ERA5 database from Copernicus (96). Evaporation is known to cause desiccation stress, and solar radiation has been shown to affect mortality and development under desiccation stress in gastropods (97, 98).

Multicollinearity is very common when working with geographical/environmental variables, so we calculated the variance inflation factor (VIF) for the geographical/environmental variables used in this study in order to remove variables that correlate with each other (99). Environmental and geographical variables were sequentially removed based on the highest VIF, until the VIF number was lower than five. A linear regression analysis, with the three variables with VIF < 5 (altitude, longitude and evaporation, the last one based on ERA5) was performed (Table S1 and Table S2A).

### RNA-seq and small RNA-seq experiments

#### Fly strains

To choose which strains to perform RNA-seq and small RNA-seq experiments on, we repeated the desiccation survival experiments with five of the most sensitive and five of the most tolerant strains according to the LT_50_ distribution (Figure S1). 3-4 replicates (except for one strain with 2 replicates) of 14-20 flies each (except for 3 replicates with 8-12 flies) for desiccation conditions, and 3-4 replicates (except for 4 strains with 1-2 replicates) of 10-20 flies (except 3 replicas that had 8-9 flies) were performed. We confirmed that tolerant strains had a higher LT_50_ compared with sensitive strains. Although the LT_50_ range was different between the two experiments (12 to 30 hours *vs* 16 to 31 hours), this was consistent with differences in humidity between the two experiments (13.65% vs 19.47%) (Table S2A-B, and Figures S1A-B). Three tolerant, GIM-012, GIM-024, and COR-023, and three sensitive TOM-08, LUN-07, and MUN-013 strains were chosen for RNA- and small RNA-sequencing (Tables S2C-D and S2D). The strains chosen had a high degree of inbreeding and a low degree of variation among biological replicates in the LT_50_ assays (Table S1-2).

#### RNA extraction for qRT-PCR and RNA-sequencing

RNA was extracted using *GenElute™* Mammalian Total RNA Miniprep Kit from 30 whole female flies, four to six days old, per replicate (three replicates per strain analyzed). RNA samples were treated with DNAse I (Thermo Fisher Scientific) following manufacturer’s instructions. RNA concentration was measured using *NanoDrop* spectrophotometer (NanoDrop Technologies) and quality was assessed with Bioanalyzer.

#### RNA extraction for small RNA sequencing

Total RNA was isolated using Trizol (Thermo Fisher Scientific) from three replicates of 30 4-7 day-old female flies after desiccation stress and under control conditions. RNA samples were treated with DNAse I (Thermo Fisher Scientific) following manufacturer’s instructions. Small RNAs were obtained by gel size selection from total RNA using 15% Mini-Protean TBE-Urea Gel (Bio-Rad, #4566056). The gel was run at 300V for 50 minutes. Fragments between 17 and 30 nucleotides were carefully cut from the gel and were put in an Eppendorf with 250μL NaCl 0.5M and then were placed at 4 °C in a rotating wheel o/n. Next, the sample was transferred to Corning Costar Spin-X centrifuge tubes (Merck, CLS8162), spinned and cleaned with standard ethanol washes and eluted in DEPEC-H2O. Quality check was performed using Bioanalyzer small RNA kit. Libraries were sequenced using Illumina Next-Seq 50 bp single-end reads.

#### RT-qPCR

We checked whether the desiccation stress triggered molecular changes in the flies and confirmed that the replicates gave similar results by measuring the expression of *frost* gene which has been reported to be up-regulated in desiccation conditions (43). Primers used were: *frost* forward (F) (5’-CGATTCTTCAGCGGTCTAGG-3’) and *frost* reverse (R) (5’-CTCGGAAAC GCCAAATTTTA-3’). RT-qPCR data were normalized with *Actin 5* (*Act5*) expression and mRNA abundance of the *frost* gene was compared to that in control samples using the 2(-Delta Delta C(T)) method and Student’s *t*-test (100) (Table S13).

### Data analysis

#### RNA-seq data

Overall, we obtained 22.4 to 42M (median 31M) paired-end reads per sample (three tolerant, three sensitive strains in treated and control conditions and three replicates per strain and condition). Fastq sequence quality was first assessed using *FastQC* (v.0.11.8) (www.bioinformatics.babraham.ac.uk/projects/fastqc) (101). Adapter and quality trimming was performed using *Cutadapt* (v. 1.18) (102) with the parameters *--quality-cutoff* 20, *-a* AGATCGGAAGAGC and the *paired-end* option. Trimmed reads were then mapped to the *D. melanogaster* genome r6.15 using *STAR* (v.2.6) (103). On average, 96.3% of the reads mapped to the reference genome. Technical duplications were explored using *dupRadar* (104). Overall, we found no bias towards a high number of duplicates at low read counts, so we did not remove duplicates from the alignments. We then used *featureCounts* (v. 1.6.2) (105) for counting the number of reads mapping to genes (*reverse-stranded* parameter). Overall, 91.81% of the aligned reads were uniquely assigned to a gene feature. We used *RSeQC* (v.2.6.4) (http://rseqc.sourceforge.net/) (106) for determining junction saturation and we found all samples saturated the number of splice junctions, meaning that the sequencing depth used in the analysis was sufficient. Raw sequencing data and matrix of raw counts per gene have been deposited in NCBI’s Gene Expression Omnibus (107) and are accessible through GEO Series accession number GSE153850.

#### Transcriptogramer analysis

*Transcriptogramer* R package (v. 1.4.1) was used to perform topological analysis, differential expression (DE), and gene ontology (GO) enrichment analyses (73). *Transcriptogramer* identifies expression profiles and analyzes GO enrichment of entire genetic systems instead of individual genes. We normalized and filtered raw counts of RNA-seq reads using the functions, *fread (), calcNormFactors()* and *filterByExpr()* (count per million values (CPM) higher than 0.5) in the *R data.table* (v. 1.12.2) (https://github.com/Rdatatable/data.table/wiki) and the *edgeR package* (v 3.24.3) (108, 109). The filtering step was performed to remove those genes that were lowly expressed and thus, would not be retained in the posterior statistical analysis. Then, we analyzed the processed data using the *Transcriptogramer* pipeline to identify the differential expression of functionally associated genes (hereafter clusters). The workflow of *Transcriptogramer* requires (i) an edge list with the gene connections, which was downloaded from *STRINGdb* (v11.0) with a combined score greater or equal to 800; (ii) an ordered gene list, where genes are sorted by the probability of their products to interact with each other, which was obtained using the *Transcriptogramer* (v 1.0) for Windows (https://lief.if.ufrgs.br/pub/biosoftwares/transcriptogramer/); (iii) expression data, which in this case were the processed reads (described above) of our RNA-seq analysis; and (iv) a dictionary, for mapping proteins to gene identifiers used as expression data row-names. The name of the genes in our data were converted to *Ensembl* Peptide IDs using the *biomaRt R/Bioconductor* package (v 2.38.0) (110) to build a dictionary to map the *Ensembl* Peptide IDs to *Ensembl* Gene IDs. First, the program assigns expression values (obtained from the RNA-seq experiment) to each respective gene in the ordered gene list. Then, the average expression of neighboring genes gets assigned to each gene in the ordered gene list. In order to measure the average expression of functionally associated genes, represented by neighbor genes in the ordered gene list, we must define a sliding window centered on a given gene with a fixed radius. We initially specified three different radii (50, 80 and 125) and finally choose 125, because this gave us the highest number of statistically significant windows per cluster (73). The p-value threshold for FDR correction for the differential expression was set to 0.01.

We then checked if the clusters of genes that were differentially expressed were enriched for specific GO terms representing specific pathways. The p-value threshold for FDR correction for the GO analysis was set to 0.005 and we focused on the first 10 GO terms with the highest adjusted p-value for each cluster when interpreting the results.

We run *Transcriptogramer* with i) the six strains comparing treated and control conditions (“All DEGs”) ii) the three tolerant strains comparing treated and control conditions (“Tolerant DEGs”) iii) the three sensitive strains comparing treated and control conditions (“Sensitive DEGs”) iv) the six strains comparing tolerant and sensitive strains in basal conditions (“Basal DEGs”).

#### Protein-protein interaction (PPI) network analysis

To identify the differentially expressed hub genes, which are likely to have a greater biological impact as they show a greater number of gene interactions, we did PPI network analysis and estimated the parameters summarizing several network properties (111). Analysis was performed using *STRING* (v 11), on the previously mentioned four groups (“All DEGs”, “Tolerant DEGs”, “Sensitive DEGs”, “Basal DEGs”). We only considered the results with a minimum required interaction score of 0.8, since those are considered as strongly correlated (112). We used *experiments* and *co-expression* data as interaction sources. The hub genes were determined using the *Cytoscape* (v.3.7.1) plugin *cytoHubba*, which calculates 11 properties of PPI networks. Among these 11 properties, Maximal Clique Centrality (MCC) is considered one of the most efficient parameters to identify hub genes (77). We ranked the genes by MCC and considered as hub genes and candidates for being involved in desiccation stress response the 30% of the genes with the highest MCC values.

#### Small RNA-sequencing data

We obtained 52-83 million single-end reads per sample (36 samples in total: three tolerant, three sensitive strains in treated and control conditions and in three biological replicates). Fastq sequence quality was assessed using FastQC (v.0.11.9) (101) and the ones that passed the filters were then used for further analysis. Afterwards, the high-quality Illumina sequencing reads were filtered using BBduk (parameters: *minlen=15 qtrim=rl ktrim=r k=21 hdist=2 mink=8;* http://jgi.doe.gov/data-and-tools/bb-tools/) to remove adapters and ribosomal RNA sequences. The trimmed reads were then analyzed using SPORTS (v.1.0.) (113): an annotation pipeline designed to optimize the annotation and quantification of canonical and non-canonical small RNAs including tRNA derived fragments (tRFs) in small RNA sequencing data (https://github.com/junchaoshi/sports1.0). SPORTS (v.1.0.) sequentially maps the cleaned reads against the *D. melanogaster* reference genome (v.6.15), miRBase (114), rRNA database (collected from NCBI), GtRNAdb (115), and piRNA database (116). On average 82% of the small RNA reads mapped to the *D. melanogaster* reference genome (r6.15). In this study, we focused on the reads mapping to the GtRNAdb (mature tRNAs) database. SPORTS (v.1.0.) was also used to identify the locations of tRFs regarding whether they were derived from 5’ terminus, 3’ terminus, or 3’CCA end of the tRNAs.

Next, to identify the genes that could be targeted by the tRFs, we mapped the tRF sequences - taken from the SPORTS output file - against the full transcripts of the *D. melanogaster* r6.15 transcriptome using RNAhybrid (117) with the following parameters: (*-u 1, -m 18500, -b 1, -v 1*). Only the first 12nt of each sequence—containing the seed region—were used as query sequences (52). As Luo *et al* (2018) suggested that 5’ fragments might have different properties to 3’ or CCA fragments under stress, all three types of tsRNA fragment were analysed separately. However, since most of the tRFs were derived from 5’ terminus (63-88%), we focused on those sequences. While two minimum free energy (MFE) cut off points were explored (MFE −20 and −30), we chose the more stringent method. This MFE −30 cut off was also chosen by Seong *et al* (2019) (118). Only transcripts with at least 10 tRF reads mapped in each one of the three replicates per sample were used for further analysis.

### Differentially expressed gene location analysis

Using the *Drosophila Gene Expression Tool* (DGET), we checked the previously reported location and the level of expression of the DEGs in this study (76). DGET uses modENCODE and RNA-seq experiment data and offers information in several life stages and tissues of *D. melanogaster.* Since we were working with 4-6 day-old females, we only used data from adult, mated four-day old females. Expression data was available for head, digestive system, carcass and ovary. We considered the genes with at least high expression (RPKM > 51) based on the DGET database. The enrichment of DEGs in tissues was checked using a hypergeometric test, and the significant p-value after a Bonferroni correction was 0.001.

### Gene disruption and RNAi knockdown strains

Three of the hub genes among the ones with the highest MCC values were selected for experimental validation (*nclb, Nsun2*, and *Dbp73D*). To determine if the candidate hub genes influence desiccation tolerance, we performed functional validation using a combination of gene disruption and RNAi transgenic lines (Table S7). The effect of each gene was tested in two different backgrounds, when stocks were available. In the case of *nclb* gene, the flies generated with the ubiquitous GAL4 driver were not viable, so we crossed the RNAi lines with the 6g1HR-GAL4-6c (Hikone) driver line, which only affects the expression of the gene of interest in the midgut, Malphigian tubules and fat body, where the *nclb* gene is mostly expressed (119) (Table S7). To generate the *Nsun2* knockdowns, we crossed strains carrying the RNAi controlled by an *UAS* promoter with flies carrying a ubiquitous GAL4 driver to silence the gene (Table S7). In the case of the *Dbp73D* gene, to overcome the lethality of the pupae caused by the ubiquitous GAL4 driver, we used an Act5c-GAL4 strain regulated by the temperature sensitive repressor GAL80 (P tubP-GAL80ts), which allowed us to time the activation of the driver. For these crosses, we transferred flies from 25°C to 29°C before emerging, to activate the driver that causes the mutation. Since not all offspring of each previously mentioned cross would have inherited the UAS-RNAi construct, we separated the flies with the construct from the ones without in the F1 generation based on the phenotypic markers. In all cases, we did reciprocal crosses of the transgenic lines and the driver strains and all experiments were carried out in the F1 generation. As controls in the experiment, reciprocal crosses with the wild-type strain of each RNAi line and the corresponding drivers were generated (Table S7).

Besides the four RNAi lines, we analyzed gene disruption strains generated with a P-element transposable element insertion for *nclb* and *Nsun2*. We used the background strain in which the mutant was generated as a control in the experiment (Table S7).

We checked if the expression of the genes was different in RNAi and gene-disruption strains compared to wild-type strains by performing RT-qPCR analysis (Table S7). For *Dbp73D,* when we crossed the #108310 female with the P tubP-GAL80ts driver male we found no difference in the expression in one of the reciprocal crosses, and slightly higher expression in the other reciprocal cross (Table S7). Thus, no further experiments were performed with this RNAi line.

All the desiccation experiments were done with 4-7 day-old females, and at least three replicates per strain were performed, using 6-13 flies/replicate (Table S7). We followed the desiccation phenotyping protocol described above. Survival curves were analyzed with log-rank test using SPSS statistical software (v.26). We found differences in the survival of the strains in control conditions in only one case. One of the reciprocal crosses for *Dbp73D* (RNAi #36131 F x Gal80ts M) showed mortality under control conditions (Table S7). Still, we found that the mortality of these flies under desiccation conditions was higher than under control conditions (long rank test p-value: <0.0001; Table S7).

### Analysis of transposable element insertions

We analyzed the TEs previously annotated in our laboratory in three tolerant (GIM-024, GIM-012, COR-023) and three sensitive (LUN-07, TOM-08, MUN-013) strains (80). Genomes of these strains were sequenced using Oxford Nanopore Technologies and TEs were *de novo* annotated using REPET package (v.2.5) (120–122). We did not consider TEs smaller than 120bp as they are known to have a high false positives/negatives rates (80).

We analyzed the TE presence/absence nearby the DEGs in the “All DEGs” and “Basal DEGs” in the six genomes, while nearby “Tolerant DEGs” were analyzed in the three tolerant strains and “Sensitive DEGs” were analyzed in the three sensitive strains. We focused on the TEs located either inside genes or within 1 kb of distance to the closest gene. To determine the enrichment or depletion of the DEGs groups nearby TEs we used Chi-square analysis.

### Desiccation-related phenotypic experiments

In the experiments detailed below, 10 tolerant and 10 sensitive strains were used from the tails of the LT_50_ phenotypic distribution, except for the respiration rate measurements, which were performed with a subset of three tolerant and three sensitive strains (Figure S1). In all the experiments, we used 4-7 day-old female flies.

#### Initial water content

The initial water content measurements were performed as described in Gibbs and Matzkin (2001) with some modifications (22). Briefly, 10 replicates of 10 females from each strain were anesthetized with CO_2_ and placed into microcentrifuge tubes, put at −80 °C for a few seconds, and then had their body weight measured. The tubes were placed at 55 °C for 72 hours and their dry body weight was measured again. The initial water content was estimated as the difference between wet and dry body mass (28).

#### Water loss analysis

Four to five replicates per strain (except for strains with 2 replicates), of 5 flies each, were anesthetized on ice and their weight was measured. Flies were then transferred to vials containing 3 grams of silica for 6 hours. After this time, their weight was measured again. The silica reduces the humidity to < 20% in about three hours, so the flies were exposed to low humidity conditions (<20%) for three hours. Water loss was calculated as the difference between the initial and final weight after desiccation stress. All the initial water content and water loss measurements were done in a *Mettler Toledo AJ100* microbalance (000.1-gram accuracy).

#### Respiration rate measurements

Insect CO_2_ exchange rate was measured with a portable photosynthesis system (Li-6400XT, Li-Cor Biosciences, Lincoln, Nebraska, USA). We used a 7.5 cm diameter clear conifer chamber (LI-6400-05; 220 cm^3^ approx. volume) to measure the respiration rate (μg CO_2_ g Insect^-1^ min^-1^). The system was calibrated daily for zero water and CO_2_ concentrations, moreover, infrared gas analyzers (IRGAs) were matched before introducing the insects in the chamber (as suggested by 6400-89 Insect Respiration Chamber Manual). Groups of five females, three replicates per strain, were placed in a net container located inside the measuring chamber. The flow rate of the air was set to 150 mol s^-1^, whereas the CO_2_ concentration inside the chamber was fixed to 400 ppm, which is the same concentration found in nature. Measurements were collected every 60 seconds. We measured for 3 hours in normal humidity conditions (65 ± 5 %) and then we reduced the humidity to 20 ± 5% and measured for 3 more hours. In both conditions, only the measurements from the second hour were used for statistical analysis, because flies need at least one hour to stabilize the respiration after the introduction in the chamber (32), and once in the chamber the rates did not change between the second and the third hour. The chamber was covered with a 2×2 cm of dark paper to keep the flies in a less active state. To test if there are differences in the respiration rate of sensitive and tolerant strains, we transformed the respiration rate values using a Yeo-Johnson transformation (bestNormalize v.1.8.2 in R). We then run two GLMM models, a model without interactions (glmm1 = lmer(R_rate_t ~ Condition + Phenotype + (1ļStrain/Replicate), res, REML=F)) and a model with interactions (glmm2 = lmer(R_rate_t ~ Condition * Phenotype + (1ļStrain/Replicate), res), REML=F), where Condition and Phenotype refers to whether the strains were under control or desiccation conditions and to the sensitive or tolerant phenotype, respectively. We then compared the two models using a likelihood ratio test (LRT).

#### Extraction and analysis of cuticular hydrocarbons

To extract the cuticular hydrocarbons (CHCs) 7-10 replicates (except for one strain with 4 replicates) of five flies for each strain were used. Flies were plunged in 200 μl of *hexane* (>99% purity, Sigma Aldrich) containing 20 μl of an internal standard (tridecane, 10ng/μl) and soaked for 9 minutes. Samples were vortexed gently for one minute and then the extract was removed and placed in a conical glass insert. The samples were stored at −20 °C until the final analysis.

Gas Chromatography Mass-Spectrometry (GS-MS) analysis was performed using a gas chromatograph (GC Agilent 7890B) coupled with a quadrupole mass spectrometer (MS Agilent 5977A MSD) operating in electron ionization mode (internal ionization source; 70 eV). 2 μl of each sample were injected in the GC injection port held at 260 °C using a split ratio of 1:5. A DB-5 ms fused silica capillary column (30m x 0.250 mm; film thickness of 0.25 μm) was used for separation using helium as the carrier gas at a constant flow rate of 1.5 mL/min. The temperature program was as follows: 40°C (1 min), then it was increased with a rate of 5°C min^-1^ until 110°C, followed by 10°C min^-1^ to 300°C (held 2 min). The mass spectra were recorded from m/z 33 to 450. A C7-C30 n-alkane series (Supelco, Bellefonte, PA) under the same chromatographic conditions was injected in order to calculate the linear retention indices (LRIs). Tentative identification of compounds was based on mass spectra matching the NIST-2014/Wiley 7.0 libraries and comparing the calculated LRIs with those available from the literature (31, 69, 74, 123–125). The amount (ng/insect) of each component was calculated relative to the internal standard (tridecane, 10ng/μl). The absolute and relative (%) amount of each component was then calculated.

Descriptive statistical analyses were performed using R (v.4.0.0.) and SPSS software (v.26). PCA analysis was performed with the log transformed data. Relative % and balanced ratios of desaturated (D) and saturated (S) compounds were calculated as in Rouault *et al* (2004): (D-S)/(D+S). Because the data was not normally distributed (Shapiro-Wilk test p-value=9.229 e-11), we used nonparametric tests. The comparison of balanced ratios and amounts of specific compounds were calculated using a Wilcoxon-signed rank test (69, 125). The correlation between the hydrocarbons and the survival of the flies (LT_100_) was calculated with Spearman’s correlation test.

### Statistical analysis

All the statistical analyses were performed using R (v.3.5.2) for Mac, unless stated differently (126).

## DECLARATIONS

### Ethics approval and consent to participate

Not applicable

### Consent for publication

Not applicable

### Availability of data and materials

All data generated and analysed during this study are included in this articles and its supplementary information files. Raw sequencing RNA-seq data and matrix of raw counts per gene have been deposited in NCBI’s Gene Expression Omnibus (107) and are accessible through GEO Series accession number GSE153850. Raw small RNA sequencing data and processed data files (results of SPORTS and RNAhybrid analysis) are accessible through GEO Series accession number GSE196669.

### Authors contribution

VH and JG designed the work. VH, SGR, JSO, EA and MR performed experiments and analyzed data, GER, LG, GA and JG analyzed data. VH and JG wrote the manuscript. All authors approved the submitted version.

## ACKNOWLEDGMENTS

We thank María Bogaerts-Márquez and Santiago Radío for technical help; Marta Coronado-Zamora and Ewan D. Harney for comments on the manuscript; Omar Rota-Stabelli for his help during the stay at the *Fondazione Edmund Mach;* and Diego Morais and Rodrigo Dalmolin for their advice related to the *Transcriptogramer* package. We also thank Tong Zhou for his help with SPORTS 1.0 package. We thank DrosEU researchers for sharing the strains analyzed in this work.

## References

1. Wheeler N, Watts N. Climate Change: From Science to Practice. Curr Environ Health Rep. 2018;5(1):170–8.

2. Parmesan C. Ecological and Evolutionary Responses to Recent Climate Change. Annual Review of Ecology, Evolution, and Systematics. 2006;37(1):637–69.

3. Stott P. CLIMATE CHANGE. How climate change affects extreme weather events. Science. 2016;352(6293):1517–8.

4. Waldvogel AM, Feldmeyer B, Rolshausen G, Exposito-Alonso M, Rellstab C, Kofler R, et al. Evolutionary genomics can improve prediction of species’ responses to climate change. Evol Lett. 2020;4(1):4–18.

5. Grillakis MG. Increase in severe and extreme soil moisture droughts for Europe under climate change. Sci Total Environ. 2019;660:1245–55.

6. Schlaepfer DR, Bradford JB, Lauenroth WK, Munson SM, Tietjen B, Hall SA, et al. Climate change reduces extent of temperate drylands and intensifies drought in deep soils. Nat Commun. 2017;8:14196.

7. Edney EB. Water Balance in Land Arthropods. Springer, Berlin 1977.

8. Gibbs AG, Chippindale AK, Rose MR. Physiological mechanisms of evolved desiccation resistance in Drosophila melanogaster. J Exp Biol. 1997;200(Pt 12):1821–32.

9. Gibbs AG, Rajpurohit S. Cuticular lipids and water balance. In G. Blomquist & A. Bagnères (Eds.), Insect Hydrocarbons: Biology, Biochemistry, and Chemical Ecology: Cambridge University Press; 2010. p. 100–20.

10. Møller AP. Quantifying rapidly declining abundance of insects in Europe using a paired experimental design. Ecol Evol. 2020;10(5):2446–51.

11. Sánchez-Bayo F, Wyckhuys KAG. Worldwide decline of the entomofauna: A review of its drivers. Biological Conservation 2019. p. 8–27.

12. Kellermann V, Heerwaarden Bv. Terrestrial insects and climate change: adaptive responses in key traits. Physiological Entomology 2019. p. 99–115.

13. Forister ML, Pelton EM, Black SH. Declines in insect abundance and diversity: We know enough to act now. Conservation Science and Practice 2019.

14. Jaramillo J, Chabi-Olaye A, Kamonjo C, Jaramillo A, Vega FE, Poehling HM, et al. Thermal tolerance of the coffee berry borer Hypothenemus hampei: predictions of climate change impact on a tropical insect pest. PLoS One. 2009;4(8):e6487.

15. Harvell CD, Mitchell CE, Ward JR, Altizer S, Dobson AP, Ostfeld RS, et al. Climate warming and disease risks for terrestrial and marine biota. Science. 2002;296(5576):2158–62.

16. Robinet C, Roques A. Direct impacts of recent climate warming on insect populations. Integr Zool. 2010;5(2):132–42.

17. Laws AN, Belovsky GE. How will species respond to climate change? Examining the effects of temperature and population density on an herbivorous insect. Environ Entomol. 2010;39(2):312–9.

18. Chown SL, Sørensen JG, Terblanche JS. Water loss in insects: an environmental change perspective. J Insect Physiol. 2011;57(8):1070–84.

19. Matos M, Simões P, Fragata I, Quina AS, Kristensen TN, Santos M. Editorial: Coping With Climate Change: A Genomic Perspective on Thermal Adaptation. Front Genet. 2020;11:619441.

20. Kellermann V, Hoffmann AA, Overgaard J, Loeschcke V, Sgrò CM. Plasticity for desiccation tolerance across Drosophila species is affected by phylogeny and climate in complex ways. Proc Biol Sci. 2018;285(1874).

21. Coyne JA, Bundgaard J, Prout T. Geographic variation of tolerance to environmental stress in Drosophila pseudoobscura.. The American Naturalist 1983. p. 474–88.

22. Gibbs AG, Matzkin LM. Evolution of water balance in the genus Drosophila. J Exp Biol. 2001;204(Pt 13):2331–8.

23. Lemeunier F, David JR, Tsacas L, Ashburner M. The melanogaster species group In M. Ashburner, H. L. Carson, & J. Thompson (Eds.), The genetics and biology of Drosophila,: London and Orlando: Academic Press; 1986. p. 147–256.

24. Parsons PA. The evolutionary biology of colonizing species. Cambridge: Cambridge University Press. 1983.

25. Hoffmann AA, Hallas RJ, Dean JA, Schiffer M. Low potential for climatic stress adaptation in a rainforest Drosophila species. Science. 2003;301(5629):100–2.

26. Hoffmann AA, Hallas R, Sinclair C, Mitrovski P. Levels of variation in stress resistance in drosophila among strains, local populations, and geographic regions: patterns for desiccation, starvation, cold resistance, and associated traits. Evolution. 2001;55(8):1621–30.

27. Matzkin LM, Watts TD, Markow TA. Desiccation resistance in four Drosophila species: sex and population effects. Fly (Austin). 2007;1(5):268–73.

28. Rajpurohit S, Oliveira CC, Etges WJ, Gibbs AG. Functional genomic and phenotypic responses to desiccation in natural populations of a desert drosophilid. Mol Ecol. 2013;22(10):2698–715.

29. Rajpurohit S, Nedved O. Clinal variation in fitness related traits in tropical drosophilids of the Indian subcontinent. Journal of Thermal Biology 2013. p. 345–54.

30. Rajpurohit S, Nedved O, Gibbs AG. Meta-analysis of geographical clines in desiccation tolerance of Indian drosophilids. Comp Biochem Physiol A Mol Integr Physiol. 2013;164(2):391–8.

31. Rajpurohit S, Zhao X, Schmidt PS. A resource on latitudinal and altitudinal clines of ecologically relevant phenotypes of the Indian Drosophila. Sci Data. 2017;4:170066.

32. Rajpurohit S, Gefen E, Bergland AO, Petrov DA, Gibbs AG, Schmidt PS. Spatiotemporal dynamics and genome-wide association genome-wide association analysis of desiccation tolerance in Drosophila melanogaster. Mol Ecol. 2018;27(17):3525–40.

33. Rouault JD, Marican C, Wicker-Thomas C, Jallon JM. Relations between cuticular hydrocarbon (HC) polymorphism, resistance against desiccation and breeding temperature; a model for HC evolution in D. melanogaster and D. simulans. Genetica. 2004;120(1-3):195–212.

34. Parkash R, Rajpurohit S, Ramniwas S. Changes in body melanisation and desiccation resistance in highland vs. lowland populations of D. melanogaster. J Insect Physiol. 2008;54(6):1050–6.

35. Parkash R, Aggarwal DD. Trade-off of energy metabolites as well as body color phenotypes for starvation and desiccation resistance in montane populations of Drosophila melanogaster. Comp Biochem Physiol A Mol Integr Physiol. 2012;161(2):102–13.

36. Hales KG, Korey CA, Larracuente AM, Roberts DM. Genetics on the Fly: A Primer on the Drosophila Model System. Genetics. 2015;201(3):815–42.

37. Griffin PC, Hangartner SB, Fournier-Level A, Hoffmann AA. Genomic Trajectories to Desiccation Resistance: Convergence and Divergence Among Replicate Selected Drosophila Lines. Genetics. 2017;205(2):871–90.

38. Telonis-Scott M, Gane M, DeGaris S, Sgrò CM, Hoffmann AA. High resolution mapping of candidate alleles for desiccation resistance in Drosophila melanogaster under selection. Mol Biol Evol. 2012;29(5):1335–51.

39. Telonis-Scott M, Sgrò CM, Hoffmann AA, Griffin PC. Cross-Study Comparison Reveals Common Genomic, Network, and Functional Signatures of Desiccation Resistance in Drosophila melanogaster. Mol Biol Evol. 2016;33(4):1053–67.

40. Kang L, Aggarwal DD, Rashkovetsky E, Korol AB, Michalak P. Rapid genomic changes in Drosophila melanogaster adapting to desiccation stress in an experimental evolution system. BMC Genomics. 2016;17:233.

41. Clemson AS, Sgrò CM, Telonis-Scott M. Transcriptional profiles of plasticity for desiccation stress in Drosophila. Comp Biochem Physiol B Biochem Mol Biol. 2018;216:1–9.

42. Cannell E, Dornan AJ, Halberg KA, Terhzaz S, Dow JAT, Davies SA. The corticotropin-releasing factor-like diuretic hormone 44 (DH44) and kinin neuropeptides modulate desiccation and starvation tolerance in Drosophila melanogaster. Peptides. 2016;80:96–107.

43. Sinclair BJ, Gibbs AG, Roberts SP. Gene transcription during exposure to, and recovery from, cold and desiccation stress in Drosophila melanogaster. Insect Mol Biol. 2007;16(4):435–43.

44. Sørensen JG, Nielsen MM, Loeschcke V. Gene expression profile analysis of Drosophila melanogaster selected for resistance to environmental stressors. J Evol Biol. 2007;20(4):1624–36.

45. Terhzaz S, Cabrero P, Robben JH, Radford JC, Hudson BD, Milligan G, et al. Mechanism and function of Drosophila capa GPCR: a desiccation stress-responsive receptor with functional homology to human neuromedinU receptor. PLoS One. 2012;7(1):e29897.

46. Sun JS, Larter NK, Chahda JS, Rioux D, Gumaste A, Carlson JR. Humidity response depends on the small soluble protein Obp59a in. Elife. 2018;7.

47. Casacuberta E, González J. The impact of transposable elements in environmental adaptation. Mol Ecol. 2013;22(6):1503–17.

48. Chuong EB, Elde NC, Feschotte C. Regulatory activities of transposable elements: from conflicts to benefits. Nat Rev Genet. 2017;18(2):71–86.

49. Van’t Hof AE, Campagne P, Rigden DJ, Yung CJ, Lingley J, Quail MA, et al. The industrial melanism mutation in British peppered moths is a transposable element. Nature. 2016;534(7605):102–5.

50. Merenciano M, Ullastres A, de Cara MA, Barrón MG, González J. Multiple Independent Retroelement Insertions in the Promoter of a Stress Response Gene Have Variable Molecular and Functional Effects in Drosophila. PLoS Genet. 2016;12(8):e1006249.

51. Guio L, Barrón MG, González J. The transposable element Bari-Jheh mediates oxidative stress response in Drosophila. Mol Ecol. 2014;23(8):2020–30.

52. Karaiskos S, Naqvi AS, Swanson KE, Grigoriev A. Age-driven modulation of tRNA-derived fragments in Drosophila and their potential targets. Biol Direct. 2015;10:51.

53. Luo S, He F, Luo J, Dou S, Wang Y, Guo A, et al. Drosophila tsRNAs preferentially suppress general translation machinery via antisense pairing and participate in cellular starvation response. Nucleic Acids Res. 2018;46(10):5250–68.

54. Dou S, Wang Y, Lu J. Metazoan tsRNAs: Biogenesis, Evolution and Regulatory Functions. Noncoding RNA. 2019;5(1).

55. Liu S, Chen Y, Ren Y, Zhou J, Ren J, Lee I, et al. A tRNA-derived RNA Fragment Plays an Important Role in the Mechanism of Arsenite-induced Cellular Responses. Sci Rep. 2018;8(1):16838.

56. Hadley NF. Water Relations of Terrestrial Arthropods. Academic Press, San Diego. 1994.

57. Hoffmann AA, Harshman LG. Desiccation and starvation resistance in Drosophila: patterns of variation at the species, population and intrapopulation levels. Heredity (Edinb). 1999;83(Pt 6):637–43.

58. Chown SL. Respiratory water loss in insects. Comp Biochem Physiol A Mol Integr Physiol. 2002;133(3):791–804.

59. Gibbs AG, Fukuzato F, Matzkin LM. Evolution of water conservation mechanisms in Drosophila. J Exp Biol. 2003;206(Pt 7):1183–92.

60. Gibbs AG. Water balance in desert Drosophila: lessons from non-charismatic microfauna. Comp Biochem Physiol A Mol Integr Physiol. 2002;133(3):781–9.

61. Telonis-Scott M, Hoffmann AA. Isolation of a Drosophila melanogaster desiccation resistant mutant. J Insect Physiol. 2003;49(11):1013–20.

62. Telonis-Scott M, Guthridge KM, Hoffmann AA. A new set of laboratory-selected Drosophila melanogaster lines for the analysis of desiccation resistance: response to selection, physiology and correlated responses. J Exp Biol. 2006;209(Pt 10):1837–47.

63. Lehmann FO, Schützner P. The respiratory basis of locomotion in Drosophila. J Insect Physiol. 2010;56(5):543–50.

64. Jallon J, Kunesch G, Bricard L, Pennanec’h M. Incorporation of fatty acids into cuticular hydrocarbons of male and female Drosophila melanogaster. J Insect Physiol. 1997;43(12): 1111–6.

65. Gibbs AG. Water-Proofing Properties of Cuticular Lipids. American Zoologist: Oxford University Press; 1998. p. 471–82.

66. Folk DG, Han C, Bradley TJ. Water acquisition and partitioning in Drosophila melanogaster: effects of selection for desiccation-resistance. J Exp Biol. 2001;204(Pt 19):3323–31.

67. Gefen E, Marlon AJ, Gibbs AG. Selection for desiccation resistance in adult Drosophila melanogaster affects larval development and metabolite accumulation. J Exp Biol. 2006;209(Pt 17):3293–300.

68. Hoffmann AA, Parsons PA. Direct and correlated responses to selection for desiccation resistance: a comparison of *Drosophila melanogaster* and *D. simulans*. Journal of Evolutionary Biology 1993. p. 643–57.

69. Ferveur JF, Cortot J, Rihani K, Cobb M, Everaerts C. Desiccation resistance: effect of cuticular hydrocarbons and water content in. PeerJ. 2018;6:e4318.

70. Foley BR, Telonis-Scott M. Quantitative genetic analysis suggests causal association between cuticular hydrocarbon composition and desiccation survival in Drosophila melanogaster. Heredity (Edinb). 2011;106(1):68–77.

71. Kennington WJ, Gilchrist AS, Goldstein DB, Partridge L. The genetic bases of divergence in desiccation and starvation resistance among tropical and temperate populations of Drosophila melanogaster. Heredity (Edinb). 2001;87(Pt 3):363–72.

72. Kellermann V, Loeschcke V, Hoffmann AA, Kristensen TN, Fløjgaard C, David JR, et al. Phylogenetic constraints in key functional traits behind species’ climate niches: patterns of desiccation and cold resistance across 95 Drosophila species. Evolution. 2012;66(11):3377–89.

73. Morais DAA, Almeida RMC, Dalmolin RJS. Transcriptogramer: an R/Bioconductor package for transcriptional analysis based on protein-protein interaction. Bioinformatics. 2019;35(16):2875–6.

74. Dembeck LM, Böröczky K, Huang W, Schal C, Anholt RR, Mackay TF. Genetic architecture of natural variation in cuticular hydrocarbon composition in Drosophila melanogaster. Elife. 2015;4.

75. Bou Sleiman MS, Osman D, Massouras A, Hoffmann AA, Lemaitre B, Deplancke B. Genetic, molecular and physiological basis of variation in Drosophila gut immunocompetence. Nat Commun. 2015;6:7829.

76. Hu Y, Comjean A, Perrimon N, Mohr SE. The Drosophila Gene Expression Tool (DGET) for expression analyses. BMC Bioinformatics. 2017;18(1):98.

77. Chin CH, Chen SH, Wu HH, Ho CW, Ko MT, Lin CY. cytoHubba: identifying hub objects and sub-networks from complex interactome. BMC Syst Biol. 2014;8 Suppl 4:S11.

78. Villanueva-Cañas JL, Horvath V, Aguilera L, González J. Diverse families of transposable elements affect the transcriptional regulation of stress-response genes in Drosophila melanogaster. Nucleic Acids Res. 2019;47(13):6842–57.

79. Ullastres A, Merenciano M, González J. Regulatory regions in natural transposable element insertions drive interindividual differences in response to immune challenges in Drosophila. Genome Biol. 2021;22(1):265.

80. Rech GE, Radio S, Guirao-Rico S, Aguilera L, Horvath V, Green L, et al. Population-scale long-read sequencing uncovers transposable elements contributing to gene expression variation and associated with adaptive signatures in Drosophila melanogaster. bioRxiv 2021.

81. Gáliková M, Dircksen H, Nässel DR. The thirsty fly: Ion transport peptide (ITP) is a novel endocrine regulator of water homeostasis in Drosophila. PLoS Genet. 2018;14(8):e1007618.

82. Lighton JR. Discontinuous gas exchange in insects. Annu Rev Entomol. 1996;41:309–24.

83. Zachariassen KE. The water conserving physiological compromise of desert insects European Journal of Entomology; 1996. p. 359–67.

84. Addo-Bediako A, Chown SL, Gaston KJ. Revisiting water loss in insects: a large scale view. J Insect Physiol. 2001;47(12):1377–88.

85. Swanson RL, Baumgardner CA, Geer BW. Very Long-Chain Fatty Acids Change the Ethanol Tolerance of Drosophila melanogaster Larvae. The Journal of Nutrition. 1995;125(3):553–64.

86. Gkatza NA, Castro C, Harvey RF, Heiß M, Popis MC, Blanco S, et al. Cytosine-5 RNA methylation links protein synthesis to cell metabolism. PLoS Biol. 2019;17(6):e3000297.

87. Kant P, Kant S, Gordon M, Shaked R, Barak S. STRESS RESPONSE SUPPRESSOR1 and STRESS RESPONSE SUPPRESSOR2, two DEAD-box RNA helicases that attenuate Arabidopsis responses to multiple abiotic stresses. Plant Physiol. 2007;145(3):814–30.

88. Genenncher B, Durdevic Z, Hanna K, Zinkl D, Mobin MB, Senturk N, et al. Mutations in Cytosine-5 tRNA Methyltransferases Impact Mobile Element Expression and Genome Stability at Specific DNA Repeats. Cell Rep. 2018;22(7):1861–74.

89. Rossi-Stacconi MV, Kaur RP, Mazzoni V, Ometto L, Grassi A, Gottardello A, et al. Multiple lines of evidence for reproductive winter diapause in the invasive pest Drosophila suzukii: useful clues for control strategies. Journal of Pest Science. 2016;89:689–700.

90. Ometto L, Cestaro A, Ramasamy S, Grassi A, Revadi S, Siozios S, et al. Linking genomics and ecology to investigate the complex evolution of an invasive Drosophila pest. Genome Biol Evol. 2013;5(4):745–57.

91. Rubel F, Kottek M. Observed and projected climate shifts 1901-2100 depicted by world maps of the K ö ppen-Geiger climate classif cation Meteorologische Zeitschrift: Gebr Ü der Borntraeger; 2010. p. 135–41.

92. Johnson RM, Dahlgren L, Siegfried BD, Ellis MD. Acaricide, fungicide and drug interactions in honey bees (Apis mellifera). PLoS One. 2013;8(1):e54092.

93. Fick SE, Hijmans RJ. WorldClim 2: new 1-km spatial resolution climate surfaces for global land areas. International Journal of Climatology 2017. p. 4302–15.

94. Copernicus CCS, (C3S). ERA5: Fifth generation of ECMWF atmospheric reanalyses of the global climate. Copernicus Climate Change Service Climate Data Store (CDS); 2017.

95. Hijmans RJ, Jakob vE. raster: Geographic analysis and modeling with raster data. R package version 2.0-12. http://CRAN.R-project.org/package=raster. 2012.

96. Bogaerts-Márquez M, Guirao-Rico S, Gautier M, González J. Temperature, rainfall and wind variables underlie environmental adaptation in natural populations of Drosophila melanogaster. Mol Ecol. 2021;30(4):938–54.

97. Przeslawski R. Combined effects of solar radiation and desiccation on the mortality and development of encapsulated embryos of rocky shore gastropods Marine Ecology Progress Series 2005. p. 169–77.

98. Strauch O, Oestergraard J, Hollmer S, Ehlers R-U. Genetic improvement of the desiccation tolerance of the entomopathogenic nematode *Heterorhabditis bacteriophora* through selective breeding Biological Control; 2004.

99. Rogerson PA. Statistical methods for geography. UK: Sage: London; 2001.

100. Livak KJ, Schmittgen TD. Analysis of relative gene expression data using real-time quantitative PCR and the 2(-Delta Delta C(T)) Method. Methods. 2001;25(4):402–8.

101. S. A. FastQC: A Quality Control Tool for High Throughput Sequence Data [Online]. Available online at: http://www.bioinformatics.babraham.ac.uk/projects/fastqc/.

102. Martin M. Cutadapt removes adapter sequences from high-throughput sequencing reads. 2011. 2011;17(1):3.

103. Dobin A, Davis CA, Schlesinger F, Drenkow J, Zaleski C, Jha S, et al. STAR: ultrafast universal RNA-seq aligner. Bioinformatics (Oxford, England). 2013;29(1):15–21.

104. Sayols S, Scherzinger D, Klein H. dupRadar: a Bioconductor package for the assessment of PCR artifacts in RNA-Seq data. BMC Bioinformatics. 2016;17(1):428.

105. Liao Y, Smyth GK, Shi W. featureCounts: an efficient general purpose program for assigning sequence reads to genomic features. Bioinformatics. 2014;30(7):923–30.

106. Wang L, Wang S, Li W. RSeQC: quality control of RNA-seq experiments. Bioinformatics. 2012;28(16):2184–5.

107. Edgar R, Domrachev M, Lash AE. Gene Expression Omnibus: NCBI gene expression and hybridization array data repository. Nucleic Acids Res. 2002;30(1):207–10.

108. Robinson MD, McCarthy DJ, Smyth GK. edgeR: a Bioconductor package for differential expression analysis of digital gene expression data. Bioinformatics. 2010;26(1): 139–40.

109. McCarthy DJ, Chen Y, Smyth GK. Differential expression analysis of multifactor RNA-Seq experiments with respect to biological variation. Nucleic Acids Res. 2012;40(10):4288–97.

110. Durinck S, Spellman PT, Birney E, Huber W. Mapping identifiers for the integration of genomic datasets with the R/Bioconductor package biomaRt. Nat Protoc. 2009;4(8): 1184–91.

111. Szklarczyk D, Gable AL, Lyon D, Junge A, Wyder S, Huerta-Cepas J, et al. STRING v11: protein-protein association networks with increased coverage, supporting functional discovery in genome-wide experimental datasets. Nucleic Acids Res. 2019;47(D1):D607–D13.

112. Zhang G, Zhang W. Protein-protein interaction network analysis of insecticide resistance molecular mechanism in Drosophila melanogaster. Arch Insect Biochem Physiol. 2019;100(1):e21523.

113. Shi J, Ko EA, Sanders KM, Chen Q, Zhou T. SPORTS1.0: A Tool for Annotating and Profiling Non-coding RNAs Optimized for rRNA- and tRNA-derived Small RNAs. Genomics Proteomics Bioinformatics. 2018;16(2):144–51.

114. Kozomara A, Griffiths-Jones S. miRBase: annotating high confidence microRNAs using deep sequencing data. Nucleic Acids Res. 2014;42(Database issue):D68–73.

115. Chan PP, Lowe TM. GtRNAdb 2.0: an expanded database of transfer RNA genes identified in complete and draft genomes. Nucleic Acids Res. 2016;44(D1):D184–9.

116. Zhang P, Si X, Skogerbø G, Wang J, Cui D, Li Y, et al. piRBase: a web resource assisting piRNA functional study. Database (Oxford). 2014;2014:bau110.

117. Krüger J, Rehmsmeier M. RNAhybrid: microRNA target prediction easy, fast and flexible. Nucleic Acids Res. 2006;34(Web Server issue):W451–4.

118. Seong KM, Coates BS, Pittendrigh BR. Impacts of Sub-lethal DDT Exposures on microRNA and Putative Target Transcript Expression in DDT Resistant and Susceptible. Front Genet. 2019;10:45.

119. Chung H, Bogwitz MR, McCart C, Andrianopoulos A, Ffrench-Constant RH, Batterham P, et al. Cis-regulatory elements in the Accord retrotransposon result in tissue-specific expression of the Drosophila melanogaster insecticide resistance gene Cyp6g1. Genetics. 2007;175(3): 1071–7.

120. Flutre T, Duprat E, Feuillet C, Quesneville H. Considering transposable element diversification in de novo annotation approaches. PLoS One. 2011;6(1):e16526.

121. Quesneville H, Bergman CM, Andrieu O, Autard D, Nouaud D, Ashburner M, et al. Combined evidence annotation of transposable elements in genome sequences. PLoS Comput Biol. 2005;1(2):166–75.

122. Hoede C, Arnoux S, Moisset M, Chaumier T, Inizan O, Jamilloux V, et al. PASTEC: an automatic transposable element classification tool. PLoS One. 2014;9(5):e91929.

123. Everaerts C, Farine JP, Cobb M, Ferveur JF. Drosophila cuticular hydrocarbons revisited: mating status alters cuticular profiles. PLoS One. 2010;5(3):e9607.

124. Flaven-Pouchon J, Farine JP, Ewer J, Ferveur JF. Regulation of cuticular hydrocarbon profile maturation by Drosophila tanning hormone, bursicon, and its interaction with desaturase activity. Insect Biochem Mol Biol. 2016;79:87–96.

125. Stinziano JR, Sové RJ, Rundle HD, Sinclair BJ. Rapid desiccation hardening changes the cuticular hydrocarbon profile of Drosophila melanogaster. Comp Biochem Physiol A Mol Integr Physiol. 2015;180:38–42.

126. Team RC. R: A language and environment for statistical computing. R Foundation for Statistical Computing, Vienna, Austria. Available online at https://www.R-project.org/. 2018.

127. Zellner BdA, Carlo B, Dugo P, Rubiolo P, Dugo G, Mondello L. Linear retention indices in gas chromatography analysis: a review. Flavour and Fragrance Journal: Wiley InterScience; 2008. p. 297–314.

